# Oxytocin release modulates acute neuroinflammation and improves brain development after pediatric traumatic brain injury

**DOI:** 10.1101/2025.06.02.652172

**Authors:** Marit Knoop, Ece Trak, Marie-Laure Possovre, Yohan van de Looij, Gabriel Schirmbeck, Kelly Ceyzériat, Jean-Luc Pitetti, Eduardo Sanches, Stefano Musardo, Philippe Millet, Stergios Tsartsalis, Benjamin B. Tournier, Camilla Bellone, Stéphane V. Sizonenko, Alice Jacquens, Olivier Baud

**Author notes:** These authors contributed equally (Alice Jacquens, Olivier Baud). Corresponding author Correspondence is invited to Olivier Baud. Contributions MK developed experimental methods, performed data collection and analysis, and wrote the manuscript. ET performed data collection and analysis. YL performed and analyzed fUS and MRI imaging experiments. JLP performed the c-casp3/Iba1/NeuN immunohistochemistry experiment. KC and PM analyzed the TSPO experiment. BBT and PM provided materials for the TSPO experiment. SM performed and analyzed the electrophysiology experiment. ST contributed to data analysis of RNA Seq experiments. GS and ES contributed to development of experimental methods and to data collection. MLP and AJ contributed to development of experimental methods and to supplementary data collection and analysis. All authors edited the manuscript. AJ and OB acquired funding and supervised the project.

## Abstract

Pediatric traumatic brain injury (TBI) is a leading cause of death and disability early in life in infants, and its neurodevelopmental consequences cannot currently be effectively treated. Since TBI is associated with neuroinflammation, modulation of the post-injury neuroinflammatory response is a promising strategy. Oxytocin is thought to have anti-inflammatory properties and appears to play a role in clinical interventions that improve brain development in neonates. However, the underlying mechanisms remain unclear, as does its applicability to acute brain injury. Here we investigate the effects of chemogenetic modulation of endogenous oxytocin on acute neuroinflammation and on long-term brain development after TBI in postnatal day 7 (P7) male mice. We show that oxytocin release attenuates the acute neuroinflammatory response to TBI 24 hours after injury, by reducing the expression of immune- and inflammation-related genes in astrocytes and promoting gene pathways for brain repair and development in microglia. In the long term, oxytocin exposure ameliorates subcortical and cortical white matter damage after TBI, prevents hyperactivity and loss of social behavior, and restores TBI-induced alterations in resting-state functional connectivity of the isocortex. These findings enhance our understanding of the modulation of neuroinflammation and its long-term effects and support intervention related to endogenous oxytocin release as a promising neuroprotective strategy in pediatric TBI.

## 1. Introduction

Pediatric traumatic brain injury (TBI) is a leading cause of death and disability in children aged 0 to 4 years^1,2^. In this age group, TBI annually affects 1.035 in 100.000 children, and is caused by falls, motor vehicle injuries or physical abuse^2^. It is heavily under-researched compared to adult TBI, despite the vast differences in injury mechanisms and effects on the brain^3^. The developing brain is particularly vulnerable to physical trauma. It has fewer tools to counteract the effects of trauma due to an underdeveloped immune system^4^, where early-life microglia are more implicated in functions related to brain development rather than immune processes^5^. Simultaneously, traumatic injury can interrupt key maturational processes taking place in the immature brain, such as myelination, synaptogenesis and neural connectivity. As a result, the effects caused by TBI in the developing brain can have a long-lasting impact on a child’s functioning in all facets of life^6,7^.

Successful clinical treatments for the pediatric population must have a proven biological basis, be as comfortable as possible for the child, and easy to implement in the clinic. However, current treatments for pediatric TBI are invasive, minimally effective, and not tailored to the pediatric population (reviewed in Figaji, 2023^8^). A novel approach on preclinical TBI research focuses on the post-injury neuroinflammatory response as therapeutic target, because modulation of microglia and astrocytes phenotypes has shown promise to influence disease outcome in brain injury models^9^. Despite promising preclinical findings, translation to clinical improvements has been largely unsuccessful in pediatric TBI. This is nourished by 3 main methodological challenges: 1) TBI is often investigated in adult animal models, 2) the type of TBI predominantly studied in animals – controlled penetrating injury – is notably different from the diffuse non-penetrating type of TBI that is characteristic in children^10^, and 3) preclinical treatments can be too complex to form a realistic option for clinical translation^11^.

One treatment option with strong clinical potential is oxytocin, a nonapeptide secreted by hypothalamic neurons, that project to all major regions of the brain – including the limbic and cortical areas^12^. Historically known for its social properties, oxytocin is gaining interest for its importance in the postnatal period and its association to brain development^13^. Some studies also support a link between oxytocin and modulation of microglial cells in brain injury models different from TBI^14–16^. Initial results associate clinical interventions that indirectly increase oxytocin levels in the newborn – such as skin-to-skin exposure^17,18^ or music therapy^19,20^ – to improved neurodevelopment of the child^21–23^. However, the causal mechanism underlying this association remains unclear. Moreover, these studies were mostly performed in preterm infants, with a varying spectrum of brain injuries^24^. Therapeutic effects of neuroinflammatory modulation have proven highly specific for the type of injury^25^ and to timeline post-injury^26,27^, so if and how oxytocin could affect inflammation specifically in pediatric TBI remains to be explored.

In this study, we assessed endogenous oxytocin release as a treatment for pediatric TBI in newborn male mice. Using a clinically-relevant model of pediatric TBI and a chemogenetic tool to manipulate oxytocinergic neuron activity, we investigated the effects of increased oxytocin release on glial reactivity following pediatric TBI, and on long-term readouts focused on myelination and white matter microstructure, functional brain connectivity and behavior. We found that only a few sessions of oxytocinergic neurons burst firing within the 4 days following TBI are sufficient to mitigate acute-phase neuroinflammation and to exert long-term positive effects on brain development.

## 2. Results

### 2.1 Traumatic brain injury causes acute neural damage and neuroinflammation in P7 male mice

Male OXT-Hm3Dq DREADD mice with a C57BL/6N background were subjected to a closed-head accelerated weight-drop model of pediatric TBI at postnatal day 7 (P7) (Fig 1a). This created cerebrovascular tissue rupture around the cortical impact site as soon as 4h post-injury, that was still present 3 days later (Fig 1b,c). Cerebrovascular damage was associated with blood accumulation in the ipsilateral subcortical white matter (Fig 1d), and increased cleaved caspase-3 apoptotic cell death in both the ipsilateral subcortical white matter and the ipsilateral cortex (located predominantly in cell layers V/VI) at 24h post-injury (Fig 1e-g). These effects are indicative of the primary injury phase of TBI. The TBI impact also induced an acute neuroinflammatory response in the ipsilateral hemisphere 24h post-injury. TBI mice showed increased CD68 phagocytic activity in the primary somatosensory cortex (Fig 1h,i) and subcortical white matter (Fig 1j,k). The increase in CD68 signal 24h after TBI was higher in the ipsilateral cortex compared to white matter (Fig 1l), reflecting a faster phagocytic response in the direct impact region rather than peripheral regions at this acute post-injury timepoint. Consistent with increased density and phagocytic activity of CD68-positive cells, a significant upregulation of microglia genes encoding for inflammatory cytokines was observed in the ipsilateral hemisphere (Fig 1m-q). The impact of the mechanical injury showed intra-group variability, explained by the immaturity of the P7 skull and dissipation of the mechanical impact – as is known for closed-head weight drop models of TBI^10^.

**Fig. 1:**
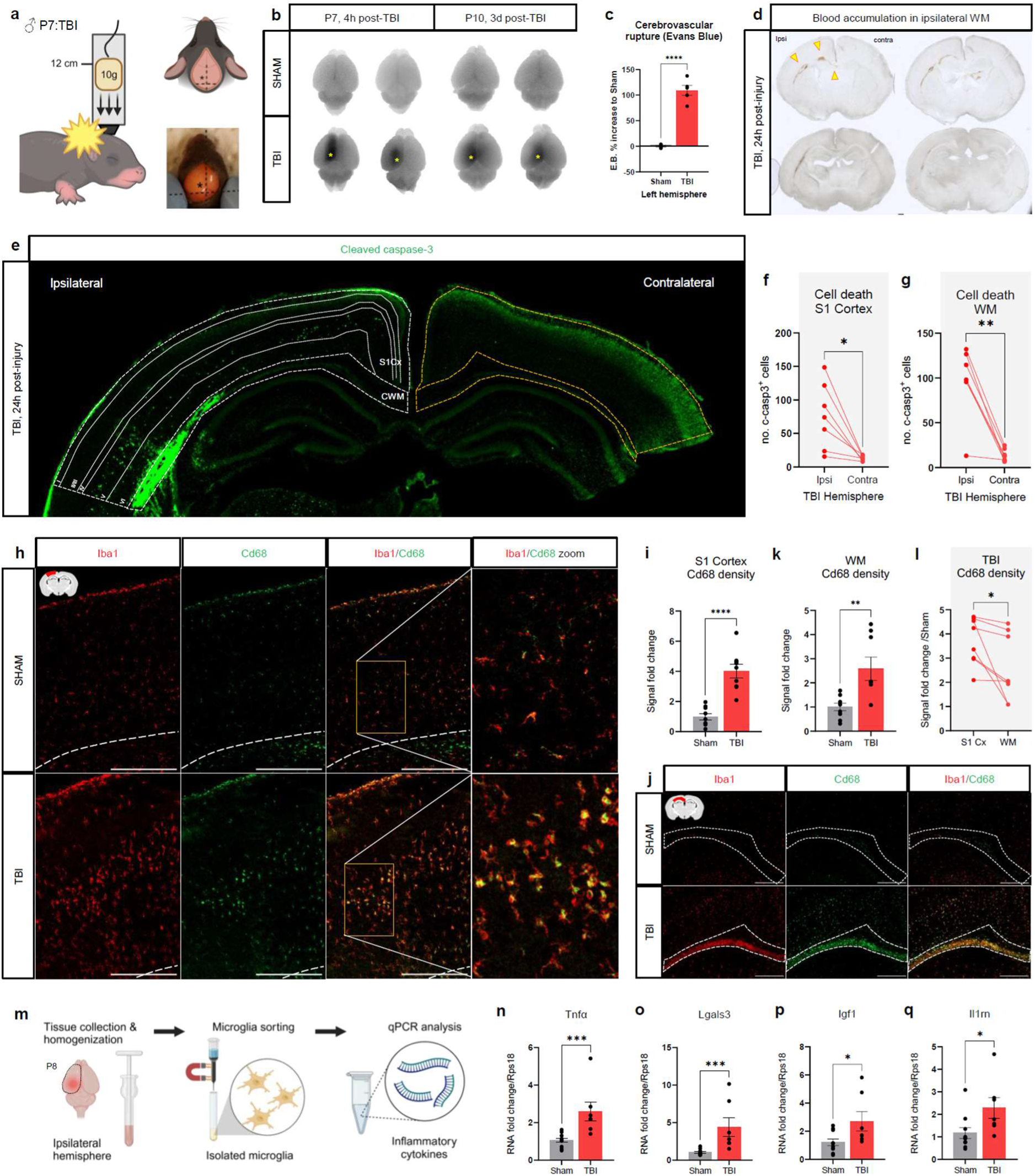
Closed-head weight drop model of pediatric TBI causes acute neural damage and neuroinflammation in P7 male mice. **a)** Graphical representation of the closed-head accelerated weight-drop model of TBI in P7 male mice, where a 10g weight is dropped from 12 cm distance to create mechanical impact directly on the exposed skull. Asterisk represents placement of impact with the dropping weight. **b)** Representative images of Evans Blue assay showing cerebrovascular rupture in the ipsilateral cortex 4 hours and 3 days after TBI injury. Yellow asterisks indicate aimed placement of the TBI dropping weight. **c)** Quantification of Evans Blue signal showing a substantial increase in cerebrovascular damage in TBI mice compared to sham. **d)** Representative images of cerebrovascular spillage into the brain 24h post-TBI. Blood accumulation is found around the ipsilateral subcortical white matter (arrows) at multiple bregma levels. **e)** Representative micrograph of cleaved-caspase 3 (green) immunohistochemistry of apoptotic cell death in the ipsilateral cortex and white matter 24 hours post-TBI. Cortical cell death was found mostly in cell layers V and VI. **f, g)** Quantification of cleaved-caspase 3 signal in ipsilateral versus contralateral hemisphere in TBI animals for S1 cortex **(f)** and cingulate white matter **(g)**, showing a unilateral injury effect of TBI for the ipsilateral hemisphere. **h,j)** Representative micrographs of Iba1 (red) and CD68 (green) signal in Sham and TBI groups 24h post-injury, as a representation of microglia/macrophage phagocytic activity in the ipsilateral cortex **(h)** and white matter **(j)**. Scale bar = 200 μm. **i,k)** Quantification of CD68 signal fold change showed that TBI increased phagocytic activity compared to sham, in both the ipsilateral cortex **(i)** and white matter **(k)**. **l)** The increase in CD68 phagocytosis in TBI mice was higher in the cortex than in the white matter. **m)** Graphical representation of qPCR assessment of microglia magnetically sorted from the ipsilateral hemisphere 24h post-TBI. **n-q)** Quantification of gene expression revealed an upregulation of neuroinflammation cytokine expression in microglia from TBI mice compared to sham mice, including Tnfα **(n)**, Lgals3 **(o)**, Igf1 **(p)**, and Il1rn **(q)**. In **(i)** Unpaired t-test (t(16) = 6.20, \*\*\*\**p* <.0001, Sham: *n* = 9, TBI: *n* = 9). In **(k)** Unpaired t-test (t(15) = 3.28, \*\**p* = .005, Sham: *n* = 9, TBI: *n* = 8). In **(l)** Paired t-test (t(7) = 2.66, \**p* = .032, *n* = 8). In **(n)** Mann-Whitney U test (U = 2, \*\*\**p* = .0004, Sham: *n* = 10, TBI: *n* = 7). In **(o)** Mann-Whitney U test (U = 2, \*\*\**p* = .0004, Sham: *n* = 10, TBI: *n* = 7). In **(p)** Mann-Whitney U test (U = 13, \**p* = .033, Sham: *n* = 10, TBI: *n* = 7). In **(q)** Mann-Whitney U test (U = 11, \**p* = .019, Sham: *n* = 10, TBI: *n* = 7). Bar and line graphs represent Mean ± SEM. Each circle represents an individual sample.

### 2.2 Oxytocin reduces microglia reactivity in the acute post-TBI phase

In this study, chemogenetic Cre-DREADD methodology^28^ was used to stimulate endogenous oxytocin release following pediatric TBI. Oxytocin release was achieved by injecting artificial ligand clozapine-n-oxide (CNO) into the OXT-Hm3Dq mice, which selectively activates the Hm3Dq receptors and increases firing of oxytocin neurons (Fig S1). Importantly, the CNO substance itself – in absence of functional DREADD expression – did not cause significant changes in readouts of this study (Fig S2).

Neuroinflammation and microglia reactivity are a major component of the secondary injury phase after TBI, that can exacerbate injury progression^9^. We assessed the effect of oxytocin on acute-phase inflammation post-TBI investigating cell death and microglia immunohistochemical features in P8 cortex. At this timepoint, TBI+OXT mice had received 2 CNO injections, 4h and 24h after brain insult (Fig 2a). At 24h post-injury, pediatric TBI mice showed an increase in cleaved caspase-3 (c-casp3) apoptotic cell death in the ipsilateral cortex, the location of the direct mechanical impact (Fig 2b,c). Co-labeling with NeuN showed that the majority of the c-casp3^+^ cells were neurons (60-80%; Fig S3). TBI mice had a higher Iba1^+^ microglial cell density in the ipsilateral cortex compared to sham (Fig 2b,d).

**Fig. 2:**
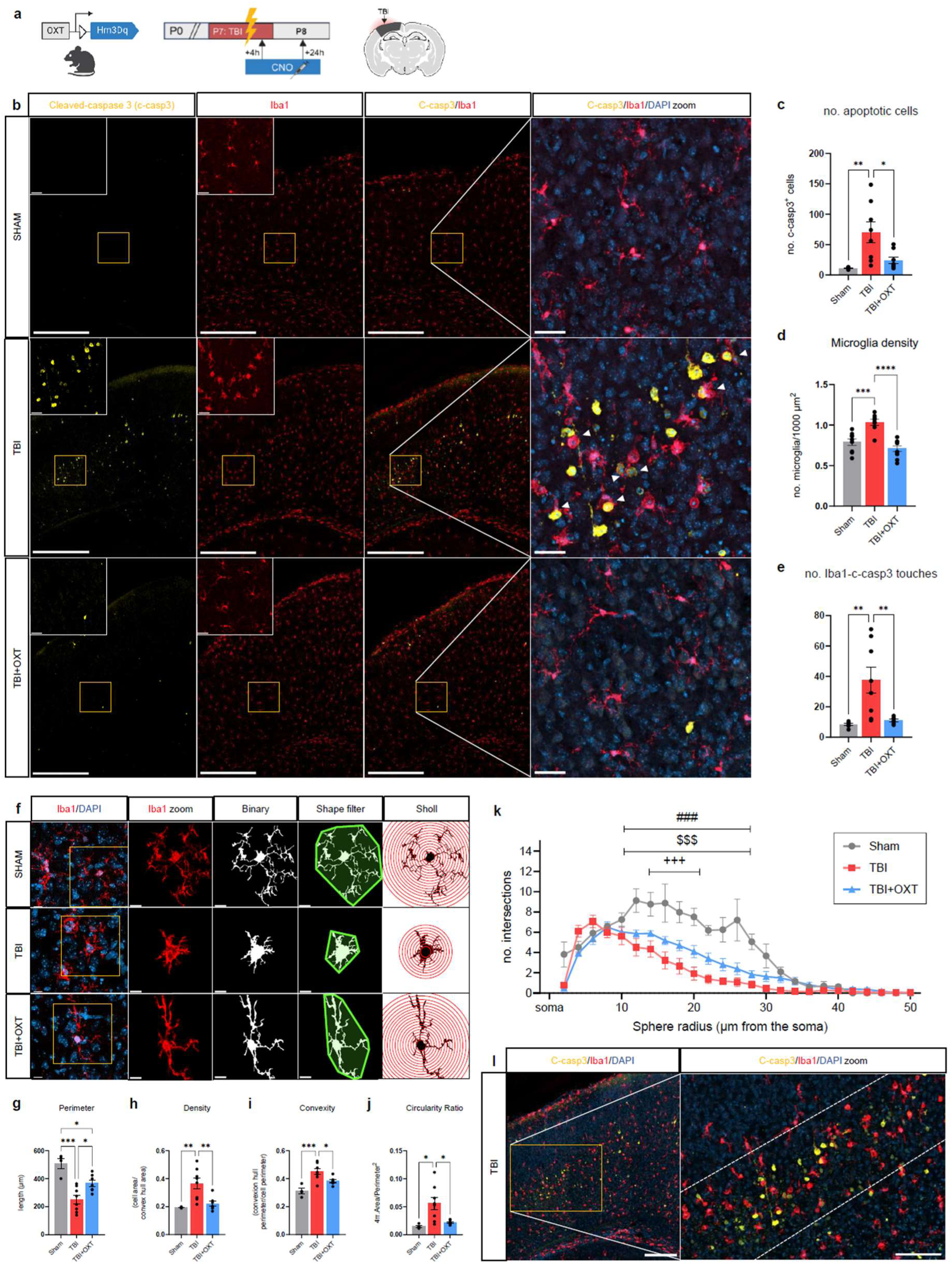
Oxytocin decreases cortical microglial reactivity 24h post-TBI. **a)** Graphical timeline of P8 immunohistochemistry experiment after P7 TBI and 2 sessions of oxytocin treatment via CNO injections, and indication of the ipsilateral somatosensory cortex ROI. **b)** Representative micrographs of apoptotic cell death (cleaved caspase-3; yellow), and microglia (Iba1; red) in the ipsilateral somatosensory cortex in Sham, TBI and TBI+OXT treatment groups. Scale bar = 200 μm (inserts: scale bar = 30 μm). Cleaved caspase-3/Iba1 co-labeling indicating apoptotic cells that are touched by a microglial cell with white arrows **(b, right column)**. **c)** Quantification of number of apoptotic cells showed an increase in TBI compared to sham, and a reverse in TBI+OXT groups. **d)** Microglia density was increased in TBI compared to sham, as was the number of Iba1-cleaved caspase-3 touches **(e)**. Both measures were reduced in the TBI+OXT group, compared to TBI **(d,e)**. **f)** Representative micrographs of individual cortical microglia in Sham, TBI and TBI+OXT groups, visualizing the processing steps for Shape filter and Sholl analysis. Scale bar = 10 μm. **g-j)** Parameters of microglia shape filter analysis revealed a more amoeboid-phenotype in TBI vs. Sham mice: decreased cell perimeter **(g)**, increased cell density **(h)**, increased cell convexity **(i)** and increased cell circularity ratio **(j)**. All shape filter parameters of cortical microglia were reversed in the TBI+OXT group compared to TBI **(g-j)**, reflecting a ramified microglia morphology similar to Sham mice. **k)** Quantification of microglia Sholl analysis showing the number of process intersections per sphere radius from the soma. ### Significant difference TBI *vs* Sham, $$$ significant difference TBI+OXT *vs* Sham, +++ significant difference TBI+OXT *vs* TBI. Sholl analysis showed that TBI microglia are less ramified than sham, and that oxytocin treatment increases the ramification degree partially. **l)** Representative micrograph of TBI mice showing an accumulation of amoeboid-shaped microglia around the cleaved caspase-3^+^ apoptotic cells in the cortex. This “amoeboid microglial line” is indicated by dashed lines. Scale bar = 100 μm (insert: scale bar = 50 μm). TBI = traumatic brain injury. CNO = clozapine n-oxide, ROI = region of interest, OXT = oxytocin. In **(c)** ANOVA (F(2,21) = 11.60, \*\**p* = .0015, Sham: *n* = 8, TBI: *n* = 8, TBI+OXT: *n* = 8). Tukey’s (Sham vs. TBI: \*\**p* = .0016, Sham vs. TBI+OXT: *p* = .65, TBI vs. TBI+OXT: \**p* = .013). In **(d)** ANOVA (F(2,24) = 19.51, \*\*\*\**p* < .0001, Sham: *n* = 9, TBI: *n* = 9, TBI+OXT: *n* = 10). Tukey’s (Sham vs. TBI: \*\*\**p* = .0005, Sham vs. TBI+OXT: *p* = .27, TBI vs. TBI+OXT: \*\*\*\**p* < .0001). In **(e)** ANOVA (F(2,19) = 9.16, \*\**p* = .0016, Sham: *n* = 8, TBI: *n* = 8, TBI+OXT: *n* = 6). Tukey’s (Sham vs. TBI: \*\**p* = .0024, Sham vs. TBI+OXT: *p* = .94, TBI vs. TBI+OXT: \*\**p* = .0098). In **(g)** ANOVA (F(2,16) = 15.97, \*\*\**p* = .0002, Sham: *n* = 4, TBI: *n* = 8, TBI+OXT: *n* = 7). Tukey’s (Sham vs. TBI: \*\*\**p* = .0001, Sham vs. TBI+OXT: \**p* = .021, TBI vs. TBI+OXT: \**p* = .023). In **(h)** ANOVA (F(2,16) = 8.93, \*\*\**p* = .0002, Sham: *n* = 4, TBI: *n* = 8, TBI+OXT: *n* = 7). Tukey’s (Sham vs. TBI: \*\**p* = .0074, Sham vs. TBI+OXT: *p* = .85, TBI vs. TBI+OXT: \*\**p* = .0075). In **(i)** ANOVA (F(2,16) = 12.59, \*\*\**p* = .0005, Sham: *n* = 4, TBI: *n* = 8, TBI+OXT: *n* = 7). Tukey’s (Sham vs. TBI: \*\*\**p* = .0004, Sham vs. TBI+OXT: *p* = .059, TBI vs. TBI+OXT: \**p* = .032). In **(j)** ANOVA (F(2,16) = 7.17, \*\**p* = .0060, Sham: *n* = 4, TBI: *n* = 8, TBI+OXT: *n* = 7). Tukey’s (Sham vs. TBI: \**p* = .016, Sham vs. TBI+OXT: *p* = .88, TBI vs. TBI+OXT: \**p* = .016). In **(k)** Two-way ANOVA (F(48,400) = 5.00, \*\*\*\**p* < .0001. Sham: *n* = 4, TBI: *n* = 8, TBI+OXT: *n* = 7). See Supplementary Data S1 for Tukey’s post-hoc statistical details. Bar graphs represent Mean ± SEM. Each circle represents an individual sample.

Single-cell analysis of microglia shape revealed a significant increase in amoeboid morphology parameters in TBI mice, including a more circular shape, higher cell density and lower number of cell ramifications (Fig 2f-k). We found an accumulation of amoeboid-shaped microglia around the c-casp3^+^ cells in TBI mice, forming an “amoeboid microglial line” around the dying cells in the ipsilateral cortex (Fig 2l). The number of Iba1^+^ / c-casp3^+^ touching cells was increased in TBI compared to sham (Fig 2e), with 57% of c-casp3^+^cells being touched by Iba1^+^ cells in TBI mice. These data suggest that in the acute phase following TBI, microglia are attracted to the dying/dead cells in the cortex and increase their presence at the impact location to start phagocytic processes. This is in line with the increase in CD68 phagocytosis immunoreactivity we found in TBI compared to sham (Fig 1h,i). TBI+OXT mice, exposed to chemogenetic oxytocin release after TBI, showed a reduction in microglia density in the ipsilateral cortex, and a reversal of amoeboid morphology 24h post-injury, compared to untreated TBI (Fig 2b,d,f-k). This was accompanied by a reduction in apoptotic cell death compared to TBI mice (Fig 2b,c). While the proportion of apoptotic neurons was unaffected by endogenous oxytocin release (Fig S3), it reduced the degree of contact between microglia and c-casp3^+^ apoptotic cells in the cortex, compared to untreated TBI (Fig 2b,e). Altogether, these data strongly suggest that oxytocin release has a significant effect on the morphologic and functional aspects of cortical microglia subjected to TBI.

Given the susceptibility of the developing white matter (WM) to any insult including TBI^3,9^, we next assessed apoptotic cell death and microglial reactivity in the ipsilateral WM 24h post-TBI (Fig 3a). TBI mice showed a substantial increase in apoptotic signal in the ipsilateral WM compared to sham group (Fig 3b,c). Similar to the cortex, the microglial response in the WM included increased cell density, more amoeboid morphology, and higher number of colocalization with c-casp3^+^ cells, compared to sham (Fig 3b,d-j). Oxytocin release reversed the increased number of microglia in the WM (Fig 3b,d), and showed a partial reversal of the amoeboid phenotype (Fig 3f-j) compared to untreated TBI mice. Notably, the apoptotic signal was unaffected by oxytocin release, as was the number of Iba1^+^ / c-casp3^+^ contacts (Fig 3b,c,e). These findings suggest that oxytocin has a potential spatial effect on cell death and acute microglial reactivity, with more evident anti-inflammatory effects in the cortex compared to the WM 24h post-injury.

**Fig. 3:**
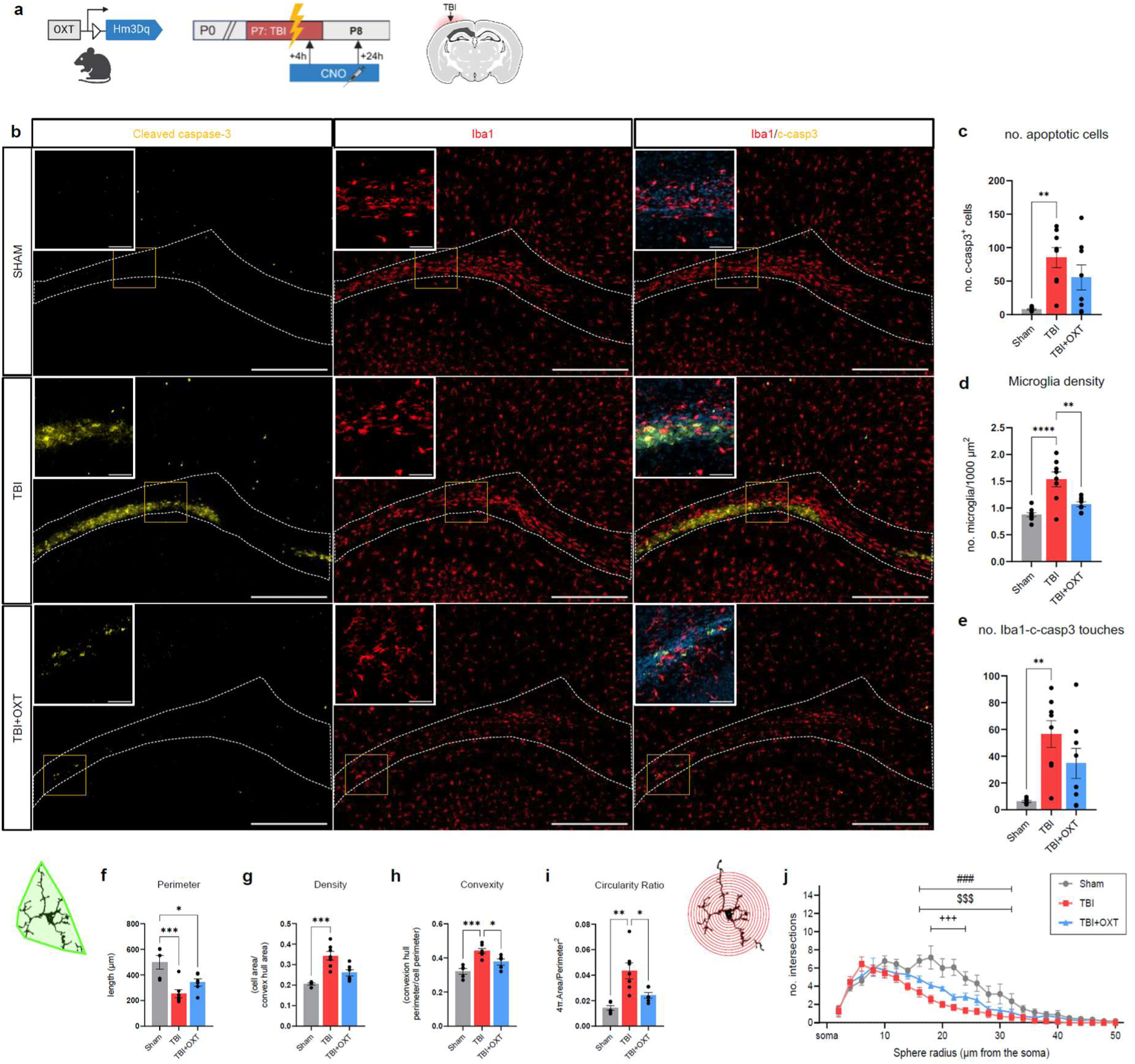
Oxytocin partially reduces white matter microglial reactivity 24h post-TBI. **a)** Graphical timeline of P8 immunohistochemistry experiment after P7 TBI and 2 sessions of oxytocin treatment via CNO injections, and graphical indication of ROI in the ipsilateral WM. **b)** Representative micrographs of apoptotic cell death (cleaved caspase-3; yellow) and microglia (Iba1; red) in the ipsilateral white matter in Sham, TBI and TBI+OXT treatment groups. Scale bar = 200 μm (inserts: 30 μm). **c)** Quantification of number of apoptotic cells showed an increase in TBI vs Sham mice, that was unaffected by the oxytocin treatment. **d)** TBI mice showed an increase in WM microglia density compared to sham. This was reduced in the TBI+OXT group compared to TBI. **e)** The number of touches between apoptotic cells and microglia was increased in TBI vs Sham, and remained high in the TBI+OXT group. **f-i)** Shape filter analysis of single microglia in the WM revealed an increase in amoeboid-morphological parameters in TBI vs Sham, including reduced cell perimeter **(f)**, increased cell density **(g)**, increased cell convexity **(h)** and increased cell circularity ratio **(i)**. TBI+OXT mice showed a partial reverse of amoeboid parameters, including a reduction in cell convexity **(h)** and cell circularity ratio **(i)**. **j)** Quantification of Sholl analysis of single microglia showed a decrease in number of intersections per sphere radius between TBI and sham groups, which was partially increased in the TBI+OXT group. ### Significant difference TBI *vs* Sham, $$$ significant difference TBI+OXT *vs* Sham, +++ significant difference TBI+OXT *vs* TBI. TBI = traumatic brain injury. CNO = clozapine n-oxide, ROI = region of interest, OXT = oxytocin. In **(c)** ANOVA (F(2,21) = 7.93, \*\**p* = .0027, Sham: *n* = 8, TBI: *n* = 8, TBI+OXT: *n* = 8). Tukey’s (Sham vs. TBI: \*\**p* = .0002, Sham vs. TBI+OXT: *p* = .06, TBI vs. TBI+OXT: *p* = .31). In **(d)** ANOVA (F(2,22) = 15.94, \*\*\*\**p* < .0001, Sham: *n* = 9, TBI: *n* = 8, TBI+OXT: *n* = 8). Tukey’s (Sham vs. TBI: \*\*\*\**p* < .0001, Sham vs. TBI+OXT: *p* = .25, TBI vs. TBI+OXT: \*\**p* = .0028). In **(e)** ANOVA (F(2,21) = 8.41, \*\**p* = .0021, Sham: *n* = 8, TBI: *n* = 8, TBI+OXT: *n* = 8). Tukey’s (Sham vs. TBI: \*\**p* = .0015, Sham vs. TBI+OXT: *p* = .076, TBI vs. TBI+OXT: *p* = .20). In **(f)** ANOVA (F(2,15) = 10.60, \*\**p* = .0014, Sham: *n* = 5, TBI: *n* = 7, TBI+OXT: *n* = 6). Tukey’s (Sham vs. TBI: \*\*\**p* = .0010, Sham vs. TBI+OXT: \**p* = .030, TBI vs. TBI+OXT: *p* = .23). In **(g)** Kruskal-Wallis test (F(3) = 13.40, \*\*\*\**p* < .0001, Sham: *n* = 5, TBI: *n* = 7, TBI+OXT: *n* = 6). Dunn’s (Sham vs. TBI: \*\*\**p* = .0008, Sham vs. TBI+OXT: *p* = .17, TBI vs. TBI+OXT: *p* = .23). In **(h)** ANOVA (F(2,14) = 15.17, \*\*\**p* = .0003, Sham: *n* = 5, TBI: *n* = 7, TBI+OXT: *n* = 5). Tukey’s (Sham vs. TBI: \*\*\**p* = .0002, Sham vs. TBI+OXT: *p* = .0582, TBI vs. TBI+OXT: \**p* = .028). In **(i)** ANOVA (F(2,16) = 10.71, \*\**p* = .0013, Sham: *n* = 5, TBI: *n* = 7, TBI+OXT: *n* = 6). Tukey’s (Sham vs. TBI: \*\**p* = .0013, Sham vs. TBI+OXT: *p* = .34, TBI vs. TBI+OXT: \**p* = .019). In **(j)** Two-way ANOVA (F(48,375) = 3.10, \*\*\*\**p* < .0001. Sham: *n* = 5, TBI: *n* = 7, TBI+OXT: *n* = 6). See Supplementary Data S2 for Tukey’s post-hoc statistical details. Bar graphs represent Mean ± SEM. Each circle represents an individual sample.

### 2.3 Oxytocin promotes gene expression of brain repair functions in microglia 24h post-TBI

We investigated the microglia transcriptome with RNA sequencing of sorted microglia from brains collected 24h post-injury at P8 (Fig 4a). Microglia cell sorting purity was verified with Expression Weighted Cell-type Enrichment analysis^29^ of the top 250 expressed genes, and confirmed the dataset was significantly enriched with only microglia cells (Fig S4a). Unsupervised PCA analysis revealed a separation of microglia transcriptome for the 3 experimental groups (Fig 4b). From the 15’933 genes sequenced, 908 genes were differentially expressed between TBI and sham microglia, of which 513 upregulated and 395 downregulated (Fig 4c,d). Gene set enrichment analysis revealed a strong upregulation of glycolytic metabolic pathways and associated pathways gluconeogenesis, response to hypoxia and HIF-1 signaling in microglia sorted from TBI-subjected animals *versus* those sorted from sham animals (TBI *vs* sham microglia) (Fig 4e). These data suggest a transcriptomic shift from homeostatic oxidative phosphorylation towards pro-inflammatory glycolysis in TBI^30,31^. Conversely, microglia sorted from TBI-subjected pups showed a downregulation of genes involved in angiogenesis and vasculogenesis compared to sham (Fig 4e), which reflects the cerebrovascular damage characteristic of 24h post-TBI^32^. We further found downregulation of gene pathways involved in brain development processes in TBI *vs* sham, including axon guidance and regulation of cell migration and differentiation (Fig 4e), suggesting that TBI-microglia contribute to impaired brain development. Many signaling pathways were downregulated in TBI *vs* sham microglia (Fig 4e): protein kinase B (ATK) signaling and the PI3K-AKT-signaling pathways, both linked with inflammation^33^, and canonical Wnt signaling, which has been linked to pro-inflammatory microglia function and hypo-myelination in the developing mouse brain^34^. Oxytocin signaling was also downregulated in TBI *vs* sham microglia (Fig 4e).

**Fig. 4:**
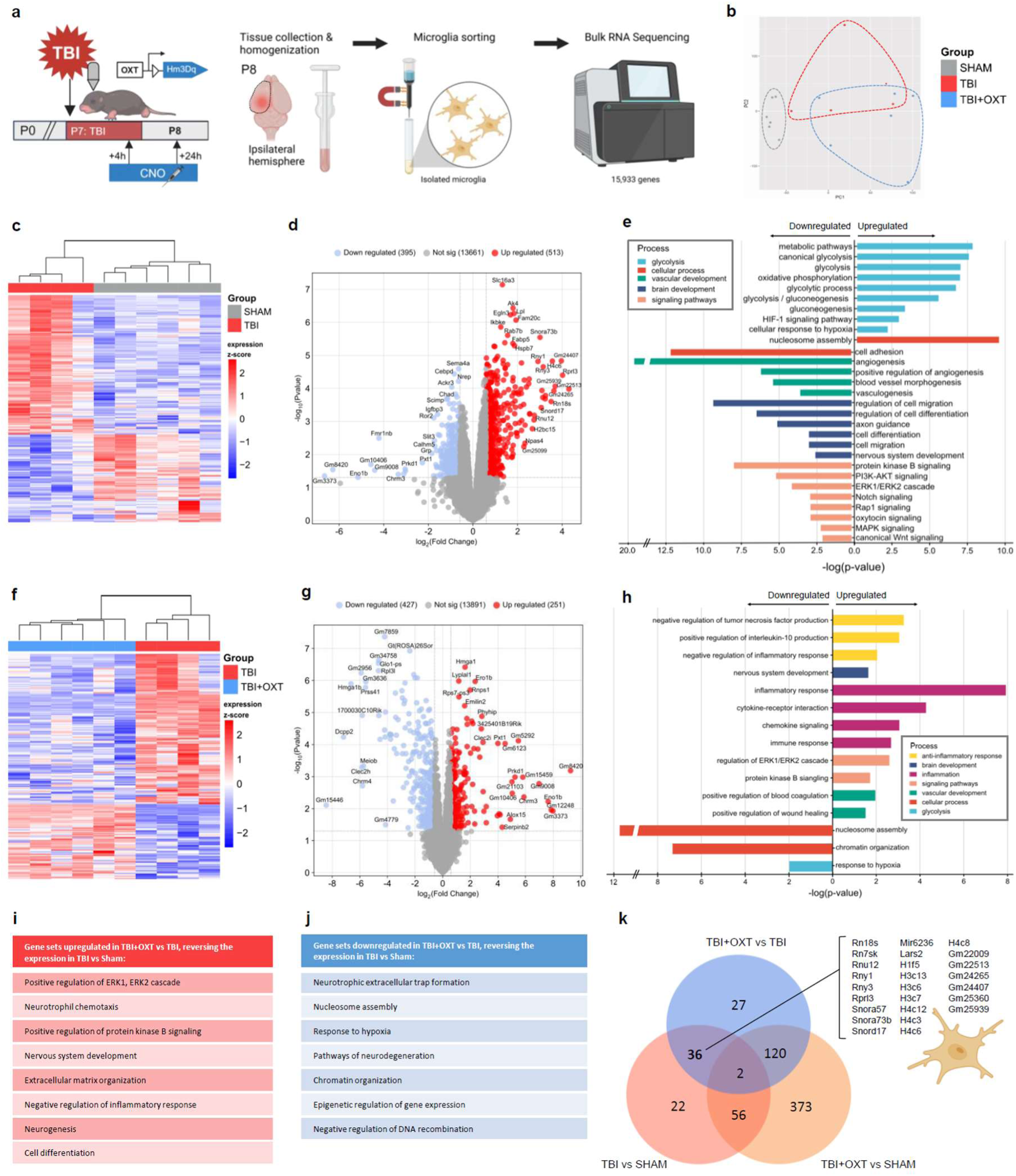
Oxytocin promotes brain repair gene pathways in microglia transcriptome 24h post-TBI. **a)** Graphical timeline of P8 microglia RNA sequencing experiment after P7 TBI and 2 sessions of oxytocin treatment via CNO injections. Microglia were magnetically sorted from ipsilateral hemispheres of P8 mice and sequenced in bulk (15’933 genes). **b)** PCA plot showing segregation of the 3 experimental groups, based on microglia transcriptomic profile. **c)** Unsupervised heatmap analysis of top deregulated genes shows separation of TBI and Sham samples into their respective treatment groups. **d)** Volcano plot showing 908 differentially expressed genes between TBI and sham microglia, including 395 downregulated (blue) and 513 upregulated (red) genes. **e)** Bar graph showing enriched gene pathways between TBI and sham microglia, assessed with gene set enrichment analysis of biological processes and KEGG repositories. TBI microglia showed an upregulation of glycolysis pathways, and a downregulation of pathways involved in vascular development, brain development and cell signaling. **f)** Unsupervised heatmap analysis of top deregulated genes between TBI+OXT and TBI microglia shows separation of TBI+OXT and TBI samples into their respective treatment groups. **g)** Volcano plot showing 678 differentially expressed genes in TBI+OXT microglia compared to TBI microglia, of which 427 downregulated (blue) and 251 upregulated (red). **h)** Bar graph of significantly enriched gene pathways in TBI+OXT microglia vs. TBI shows an upregulation of (anti-) inflammatory pathways, a rescue of vascular development and brain development pathways, and a downregulation of glycolysis and cellular processes. **i,j)** Tables highlighting gene sets upregulated **(i)** and downregulated **(j)** in TBI+OXT vs TBI, reversing the transcriptomic effect of TBI vs Sham in sorted microglia. **k)** Venn diagram of differentially expressed genes between treatment groups revealed 36 genes common genes between TBI vs. Sham and TBI+OXT vs. TBI comparisons. These genes were associated with histone and chromatin compartments. These data suggest that oxytocin treatment promotes anti-inflammatory gene expression in microglia 24h post-TBI, as well as improved expression of brain and cerebrovascular development gene pathways. TBI = traumatic brain injury. CNO = clozapine n-oxide, OXT = oxytocin. DEG = differentially expressed gene, GSEA = gene set enrichment analysis. RNA Seq analysis was performed with > 1.5 thresholds for fold change, and with FDR-corrected p-value < .05 for the DEG analysis and p < .05 for the GSEA analysis. In **(e,h)** enrichment is represented as the negative log of the p value. Upregulated pathways are positioned on the right side of the y-axis, and downregulated pathways on the left.

Oxytocin treatment affected expression of 678 microglial genes compared to untreated TBI microglia, of which 251 upregulated and 427 downregulated (Fig 4f,g). The upregulated genes in microglia sorted from animals exposed to TBI and OXT were enriched for pathways associated with anti-inflammatory functions (Fig 4h). Oxytocin also increased gene expression of vascular wound healing pathways compared to untreated TBI, which suggests improved repair of the cerebrovascular damage caused by TBI. Neurodegeneration was one of the pathways that was downregulated in TBI+OXT microglia, as were general cellular processes such as nucleosome assembly and chromatin organization (Fig 4h).

We identified several pathways whose differential upregulation in TBI *vs* sham microglia was directly reversed by endogenous oxytocin release (Fig 4i,j). It reversed the downregulation of neurogenesis, cell differentiation and nervous system development pathways, as well as signaling pathways like ERK1/ERK2 and protein kinase B in TBI (Fig 4i). Likewise, oxytocin reversed the upregulation of pathways involved in neurodegeneration and hypoxia found in TBI, as well as epigenetic pathways such as chromatin organization (Fig 4j). Notably, the epigenetic effects were the most significant of all the transcriptomic pathways that were affected by oxytocin. These data suggest that oxytocin promotes the expression of tissue repair and brain development genes in microglia 24h post-TBI. Moreover, the 36 overlapping genes that were differentially expressed by TBI (*vs* sham) and also by TBI+OXT (*vs* TBI) showed an enrichment for histone (cluster 3 and 4) and chromatin compartments (Fig 4k). These genes were all downregulated by oxytocin compared to untreated TBI, restoring their expression to Sham-equivalent levels.

### 2.4 Oxytocin differently affects the astrocyte transcriptome from microglia 24h post-TBI

Next, we aimed to map the effects of oxytocin on astrocytes. Given that astrocytes and microglia have the ability to bidirectionally influence each other^35^, we first wondered if the effects of oxytocin on microglia were direct, or indirect effects mediated by astrocyte signaling. Using RNA scope *in situ* hybridization on P5 brain sections, we found support that both microglia and astrocytes express oxytocin receptor mRNA *in-vivo*, in both cortical and white matter regions (Fig S5) – as consistent with previous studies^15,36^. This confirms that P7 microglia can be directly influenced by oxytocin – which had not been demonstrated yet *in-vivo* for young mice (reviewed in Knoop et al., 2022^13^). We then performed RNA seq on astrocytes to understand transcriptomic changes caused by TBI and oxytocin exposure, to complement the microglia results.

Astrocytes were sorted from ipsilateral hemispheres 24h post-TBI at P8 and analyzed with RNA sequencing (Fig 5a,b). Expression Weighted Cell-type Enrichment analysis^29^ of the top 250 expressed genes confirmed the cell sorting purity, showing a significantly enrichment of only astrocytes in the dataset (Fig S4b). We sequenced 17’460 genes and found 357 genes expressed differentially between TBI *vs* sham astrocytes, of which 269 upregulated and 88 downregulated (Fig 5c,d). Gene set enrichment analysis revealed a significant upregulation of inflammatory pathways in TBI astrocytes compared to sham, including a cluster of phagocytosis pathways (Fig 5e). Similarly, there was an abundant upregulation of pathways involved in the immune response, which included regulation and proliferation of cells of the adaptive immune response, and a specific activation of microglia (Fig 5e). This transcriptomic profile of TBI astrocytes was in line with the well-known astrocytic pro-inflammatory response in the acute post-injury phase^37^. Signaling pathways that were upregulated in TBI astrocytes included ERK1/ERK2 cascade, NF kappa B signaling, and MAPK and JNK pathways (Fig 5e), which are known to drive astrocyte reactivity in inflammation^38^. Downregulated gene pathways in TBI astrocytes involved synaptic functioning (Fig 5e), one of the main neuron-support functions of astrocytes in homeostasis^39^.

**Fig. 5:**
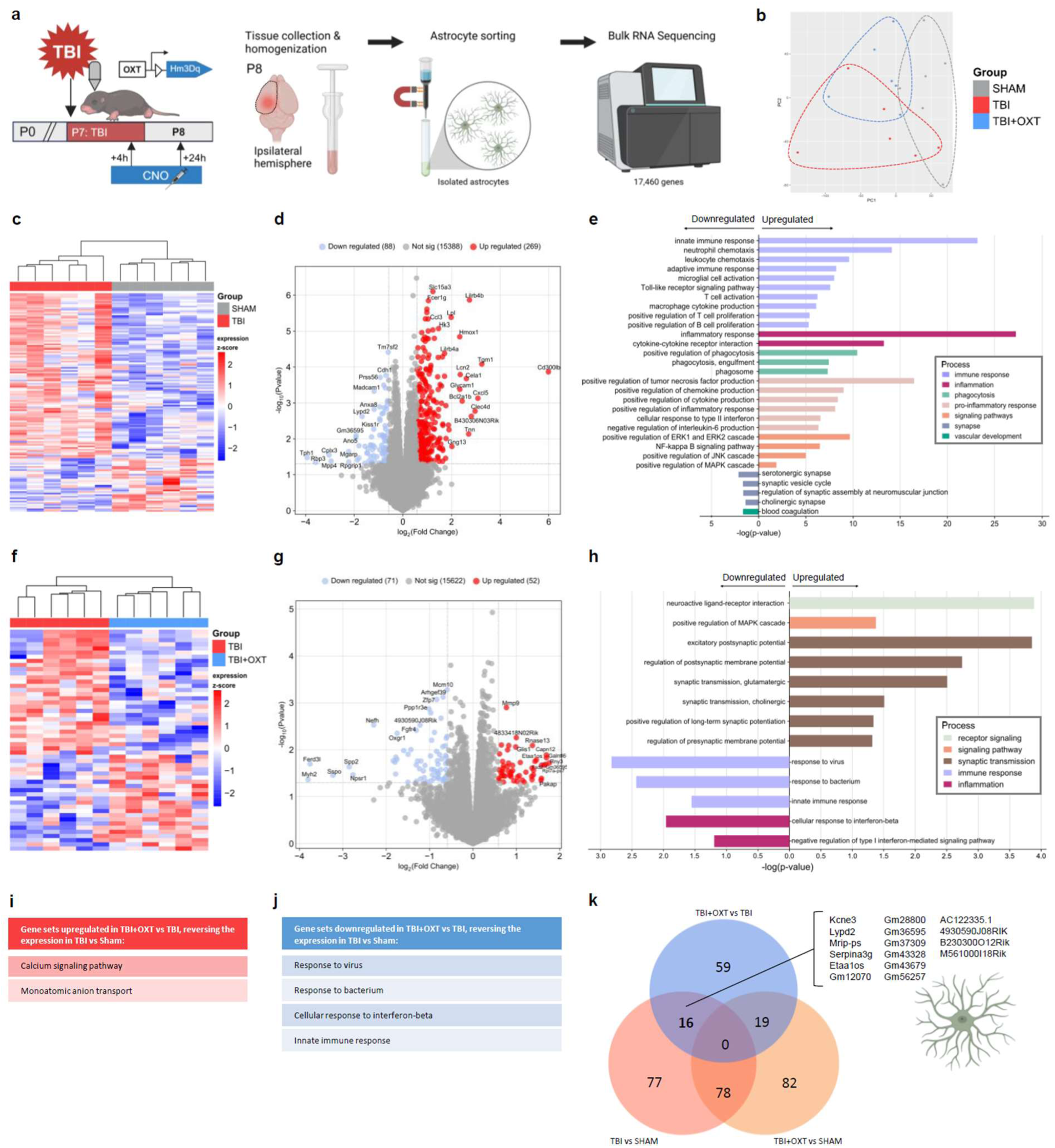
Oxytocin dampens immune and inflammation gene pathways in astrocyte transcriptome 24h post-TBI. **a)** Graphical timeline of P8 astrocyte RNA sequencing experiment after P7 TBI and 2 sessions of oxytocin treatment via CNO injections. Astrocytes were magnetically sorted from ipsilateral hemispheres of P8 mice and sequenced in bulk (17’460 genes). **b)** PCA plot showing segregation of the 3 treatment groups, based astrocyte transcriptomic profile. **c)** Unsupervised heatmap analysis of top deregulated genes shows separation of TBI and Sham samples into their respective treatment groups. **d)** Volcano plot showing 357 differentially expressed genes between TBI and sham astrocytes, including 88 downregulated (blue) and 269 upregulated (red) genes. **e)** Bar graph showing significantly enriched gene pathways between TBI and sham astrocytes, assessed with gene set enrichment analysis of Biological processes and KEGG repositories. TBI astrocytes showed a strong upregulation of immune and inflammation gene pathways, and a downregulation of synaptic pathways. **f)** Unsupervised heatmap analysis of top deregulated genes between TBI+OXT and TBI astrocytes shows separation of TBI+OXT and TBI samples into their respective treatment groups. **g)** Volcano plot showing 123 differentially expressed genes in TBI+OXT astrocytes compared to TBI astrocytes, of which 71 downregulated (blue) and 52 upregulated (red). **h)** Bar graph of significantly enriched gene pathways in TBI+OXT astrocytes vs. TBI shows an upregulation of synaptic processes and a downregulation of immune and inflammation pathways. **i,j)** Tables highlighting gene sets upregulated **(i)** and downregulated **(j)** in TBI+OXT vs TBI, reversing the transcriptomic effect of TBI vs Sham in sorted astrocytes **k)** Venn diagram of differentially expressed genes between treatment groups revealed 16 genes commonly shared between TBI vs. Sham and TBI+OXT vs. TBI comparisons. These genes were not associated with any particular processes or cellular compartments. The data suggest that oxytocin treatment reduces the pro-inflammatory and immune response in astrocytes 24h post-TBI. No effects on brain development pathways were found for astrocytes. TBI = traumatic brain injury. CNO = clozapine n-oxide, OXT = oxytocin. DEG = differentially expressed gene, GSEA = gene set enrichment analysis. RNA Seq analysis was performed with > 1.5 thresholds for fold change, and with FDR-corrected p-value < .05 for the DEG analysis and p < .05 for the GSEA analysis. In **(e,h)** enrichment is represented as the negative log of the p value. Upregulated pathways are positioned on the right side of the y-axis, and downregulated pathways on the left.

Astrocytes sorted from TBI+OXT pups had 123 genes differentially expressed compared to those sorted from untreated TBI pups, of which 52 upregulated and 71 downregulated (Fig 5f,g). TBI+OXT astrocytes showed a reversal of some upregulated pathways specifically involved in inflammation and the immune response observed in TBI *vs* sham, including response to type 1 interferon (Fig 5h). No effects of oxytocin on immune cell recruitment or microglia activation were found. The upregulated genes in TBI+OXT astrocytes were enriched for pathways associated with regulation of the MAPK cascade and synaptic transmission, including presynaptic and postsynaptic processes (Fig 5h). Unlike its effect on microglia, oxytocin did not regulate genes involved in vascular tissue recovery or brain development in TBI astrocytes.

Gene pathways increased in TBI *vs* sham and subsequently decreased by endogenous oxytocin release involved the immune response and response to virus (Fig 5j). The 16 overlapping genes between TBI+OXT (*vs* TBI) and TBI (*vs* sham) microglia were not enriched for any specific functions or cellular compartments (Fig 5k). Oxytocin did not affect brain development processes or microglial signaling in astrocytes. These data suggest that the transcriptomic effects of OXT on astrocytic acute response to TBI mostly involve a reduction of immune-associated pro-inflammatory gene pathways, and are different from the effects of oxytocin release on microglia after TBI.

### 2.5 Long-term white matter microstructural deficiencies following TBI are affected by oxytocin release

Besides acute effects of oxytocin on post-TBI neuroinflammation, we wanted to assess the long-term effects of oxytocin release on brain maturation and function following TBI. For these long-term readouts at P45, mice from the TBI+OXT treatment group received daily CNO injections between 4h post-TBI (P7) and 3d post-TBI (P10), for a total of 4 injections. We first assessed white matter development, using immunohistochemistry of myelin base protein (MBP) (Fig 6a). TBI mice had smaller corpus callosum thickness at P45, which was reversed in the TBI+OXT group (Fig 6b,c). The ipsilateral external capsule was substantially thinner in TBI mice compared to sham, and unaffected by oxytocin treatment (Fig 6d,e). We also assessed myelination of the ipsilateral S1 cortex (the “TBI impact site”) and found local effects of TBI on axonal myelination (Fig 6f-h): TBI mice showed lower MBP density and reduced myelin coverage of axons compared to sham mice. Oxytocin reversed these deficits, restoring sham-equivalent levels of ipsilateral cortical MBP content and signal intensity. Finally, coherency of cortical myelin – an inverted measure of fiber integrity^40^ – was not different between groups (Fig 6i).

**Fig. 6:**
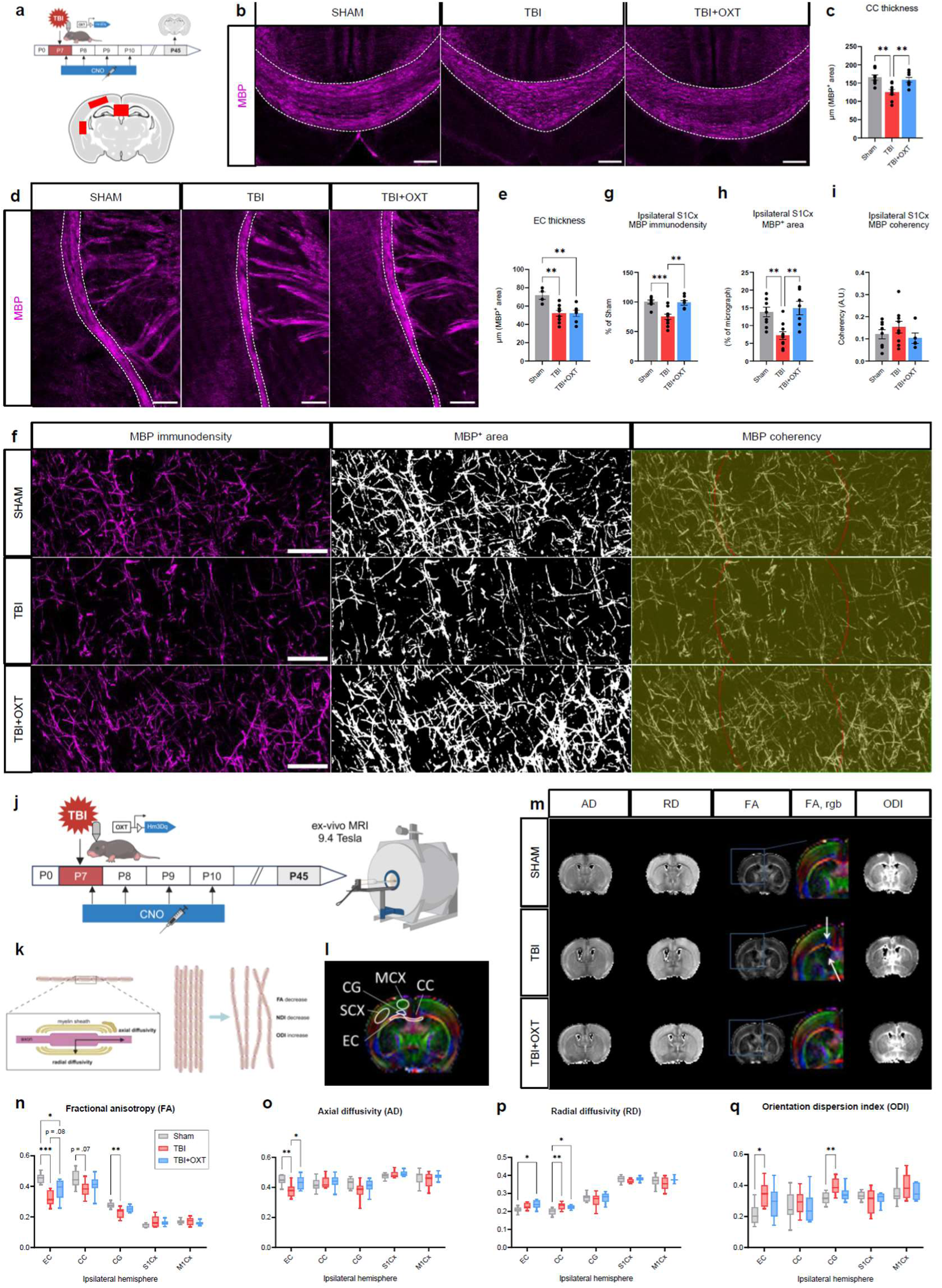
Long-term white matter microstructural deficiencies following TBI are affected by oxytocin treatment. **a)** Graphical timeline of the P45 myelin base protein (MBP) immunohistochemistry experiment, with the 3 ROIs in red. **b,d**) Representative micrographs of MBP signal (magenta) in the CC **(b)** and EC **(d)** for Sham, TBI and TBI+OXT groups. Scale bar = 100 μm. **c)** Quantification of CC thickness shows a reduction in TBI and a rescue in the TBI+OXT group. **e)** Quantification of EC thickness shows a reduction in TBI vs. Sham mice, and no effect of the oxytocin treatment. **f)** Representative micrographs of MBP immunodensity (left), MBP^+^ area (middle) and MBP coherency (right) of ipsilateral cortical white matter for Sham, TBI and TBI+OXT groups. Scale bar = 20 μm. **g-i)** Quantification of cortical WM measures shows a decrease in MBP immunodensity **(g)** and MBP^+^ area **(h)** in TBI, which were both rescued in TBI+OXT mice. MBP coherency was not affected by treatment group **(i)**. **j)** Graphical timeline of the P45 *ex-vivo* DTI-MRI experiment after P7 TBI and P7-P10 oxytocin treatment via CNO injections. **k)** Graphical representation of the DTI and NODDI parameters, illustrating the different aspects of white matter microstructure they reflect. **l)** Atlas of ipsilateral ROIs that were quantified from MRI scans: 3 WM ROIs (CC, EC, CG) and 2 gray matter ROIs (MCX and SCX). **m)** Representative MRI sections of Sham, TBI and TBI+OXT mice showing the AD, RD, FA, ODI and NDI signals in the ROIs. Arrows emphasize FA damage in WM regions of TBI mice. **n-r)** Quantification of DTI **(n-p)** and NODDI **(q)** parameters between Sham (gray), TBI (red) and TBI+OXT (blue) groups. AD **(o)** and RD **(p)** are expressed in x10^-2^ mm^2^.s^-1^. Compared to sham mice, TBI mice showed a reduction in FA **(n)** for all WM ROIs, reduced AD **(o)** in the EC, increased RD **(p)** in the EC and CC, and increased ODI **(q)** in the EC and CG. Compared to TBI mice, TBI+OXT mice showed an increased FA **(n)** in the EC, and an increased AD **(o)** in the EC. TBI = traumatic brain injury, CNO = clozapine n-oxide, CC = corpus callosum, EC = external capsule, CG = cingulum, MCX = primary motor cortex, SCX = primary somatosensory cortex, AD = axonal diffusion, RD = radial diffusivity, FA = fractional anisotropy, ODI = orientation dispersion index, NDI = neurite density index. DTI = diffusion tensor imaging, NODDI = neurite orientation dispersion and density imaging, WM = white matter, MBP = myelin base protein. In **(c)** ANOVA (F(2,22) = 9.71, \*\*\**p* = .0009, Sham: *n* = 8, TBI: *n* = 9, TBI+OXT: *n* = 8). Tukey’s (Sham vs. TBI: \*\**p* = .0014; Sham vs. TBI+OXT: *p* = .80; TBI vs. TBI+OXT: \*\**p* = .0067). In **(e)** ANOVA (F(2,17) = 7.21, \*\**p* = .0052, Sham: *n* = 4, TBI: *n* = 9, TBI+OXT: *n* = 7). Tukey’s (Sham vs. TBI: \*\**p* = .0065; Sham vs. TBI+OXT: \*\**p* = .0092; TBI vs. TBI+OXT: *p* = 1). In **(g)** ANOVA (F(2,22) = 12.96, \*\*\**p* = .0002, Sham: *n* = 8, TBI: *n* = 10, TBI+OXT: *n* = 8). Tukey’s (Sham vs. TBI: \*\*\**p* = .0005; Sham vs. TBI+OXT: *p* = .97; TBI vs. TBI+OXT: \*\**p* = .0013). In **(h)** ANOVA (F(2,23) = 9.86, \*\*\**p* = .0008, Sham: *n* = 9, TBI: *n* = 10, TBI+OXT: *n* = 7). Tukey’s (Sham vs. TBI: \*\**p* = .0041; Sham vs. TBI+OXT: *p* = .85; TBI vs. TBI+OXT: \*\**p* = .0019). In **(i)** ANOVA (F(2,19) = 0.91, *p* = .42, Sham: *n* = 8, TBI: *n* = 9, TBI+OXT: *n* = 5). In **(n-q)** See Supplementary Data S3 for statistical details. Bar graphs and box plots represent Mean ± SEM. Each circle represents an individual sample.

To further explore white matter integrity, we performed *ex-vivo* magnetic resonance diffusion tensor imaging at P45 in the corpus callosum (CC), external capsule (EC), cingulum, primary somatosensory cortex (S1Cx) and primary motor cortex (M1Cx) of the ipsilateral hemisphere (Fig 6j-m). Compared to sham, TBI mice showed reductions in fractional anisotropy in the EC, cingulum and a trend for the CC (p = .08) (Fig 6n). TBI mice had significantly lower axial diffusivity in the EC, which reflects axonal damage (Fig 6o). Increased radial diffusivity was found statistically significant in the CC in TBI mice, indicating damage to the myelin sheaths (Fig 6p). NODDI analysis of neurite microstructure further revealed increased orientation dispersion in the EC and cingulum in TBI mice compared to sham, which reflects a loss of coherence in axon orientation (Fig 6q). No effects were found of TBI on microstructural characteristics of the S1 and M1 cortical regions. TBI+OXT mice showed a significant rescue of axial diffusivity in the EC compared to TBI mice at P45 (Fig 6o). Similarly, they showed a trend to reversal of fractional anisotropy damage in the EC (p = .08) (Fig 6n). No other effects of oxytocin treatment were found on long-term MRI microstructure following TBI.

### 2.6 Oxytocin improves loss of long-term functional brain connectivity after TBI

Next, we aimed to correlate microstructural changes induced by TBI and oxytocin release with brain function. We assessed the long-term neural correlates of pediatric TBI and oxytocin exposure using functional ultrasound (fUS), an *in-vivo* imaging technique based on blood flow variation, that can be used to assess cerebrovascular integrity and calculate correlations of neural activity between brain regions (e.g., “functional connectivity”)^41^ (Fig 7a-c). fUS analysis revealed a loss of cerebral blood volume (CBV) in the ipsilateral hemisphere in P45 TBI mice compared to sham (Fig 7d,e), demonstrating that the damage induced by the traumatic impact at P7 has long-lasting effects on brain vasculature. TBI+OXT mice showed increased, sham-equivalent CBV in the ipsilateral hemisphere compared to untreated TBI (Fig 7d,e), suggesting improved long-term cerebrovascular integrity after early-life oxytocin treatment.

**Fig. 7:**
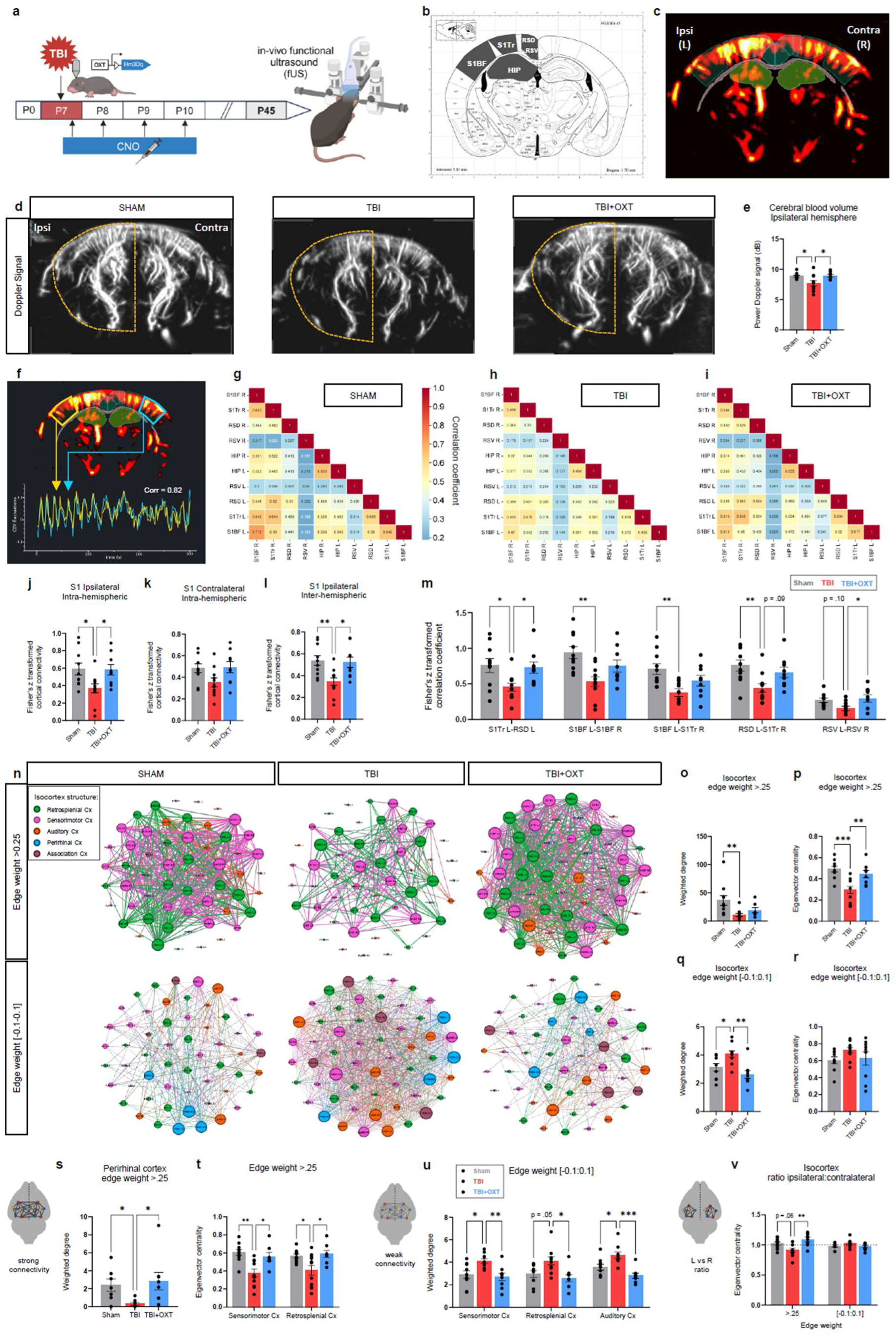
Oxytocin improves long-term functional brain connectivity after TBI. **a)** Graphical timeline of the P45 *in-vivo* resting-state fUS experiment after P7 TBI and P7-P10 oxytocin treatment via CNO injections. **b)** Visualization of the bregma plane and the 10 ROIs that were selected for correlation connectivity analysis: S1Bf, S1Tr, RSV, RSD, HIP, in the ipsilateral and contralateral hemisphere. **c)** Example of the cerebral blood volume signal of a mouse with the ROIs superimposed. Lighter colors represent higher CBV, red colors represent lower CBV volumes. **d)** Representative images of the CBV as a measure of cerebrovascular integrity in the Sham (left), TBI (middle) and TBI+OXT (right) groups, with ipsilateral hemisphere delineated with dotted line. **e)** Quantification of CBV in the ipsilateral hemisphere shows a reduction in TBI mice versus Sham, and a rescue in the TBI+OXT group. **f)** Visualization of how the functional connectivity was calculated: as a correlation coefficient between the CBV variations between 2 ROIs. This example shows high correlation (r = .82). **(g-i)** Average correlation coefficient matrices of functional connectivity between the 10 ROIs for Sham **(g)**, TBI **(h)** and TBI+OXT **(i)** groups. **j-l)** Quantification of cortical functional connectivity in the S1 regions, divided into intra-hemispheric ipsilateral **(j)**, intra-hemispheric contralateral **(k)** and inter-hemispheric **(l)** connectivity. TBI mice showed a reduction, and TBI+OXT mice a rescue, for ipsilateral intra-hemispheric connectivity **(j)**, inter-hemispheric connectivity **(l)**, but no effect was found on contralateral intra-hemispheric connectivity **(k)**. **m)** Quantification of functional connectivity between a selection of ROI-pairs for Sham (gray), TBI (red) and TBI+OXT (blue) groups. **n-w)** Network analysis of functional connectivity on fUS registrations from 212 ROIs across 3 bregma planes. **n)** Average isocortex networks of the Sham (left), TBI (middle) and TBI+OXT (right) groups, separated into strong connections (edge weight >.25; top) and weak connections (edge weight between -0.1 and 0.1; bottom). Visualized with the Fruchterman-Reingold force-directed layout algorithm. Node size represents eigenvector centrality (EC), and edge size represents weighted degree (= strength; WD). Colors represent the 5 isocortex substructures: retrosplenial cortex (green), sensorimotor cortex (magenta), auditory cortex (orange), perirhinal cortex (blue) and association cortex (mauve). **o,p)** Quantification of WD **(o)** and EC **(p)** measures for strong connections of the isocortex shows a decrease in both WD **(o)** and EC **(p)** in the TBI group versus sham, and a rescue of EC **(p)** in the TBI+OXT group. **q,r)** Quantification of WD **(q)** and EC **(r)** for weak connections of the isocortex showed an increase in TBI and a reverse in TBI+OXT for WD **(q)**. No effect on EC was found for weak isocortex connections **(r)**. **s)** Quantification of strong connectivity WD revealed a decrease in TBI and increase in TBI+OXT mice for the perirhinal cortex. **t)** Quantification of strong connectivity EC showed a decrease in TBI (red) and reversal in TBI+OXT (blue) for the sensorimotor and retrosplenial cortices. **u)** Quantification of weak connectivity WD for a selection of isocortex substructures for Sham (gray), TBI (red) and TBI+OXT (blue) groups, showed an increase in TBI and rescue in TBI+OXT for the sensorimotor, retrosplenial and auditory cortices. **v)** Quantification of the ratio of EC between ipsilateral/left and contralateral/right isocortex regions as a representation of hemispheric balance revealed a preference for the contralateral regions in TBI (red) and a restoration of equal balance in the TBI+OXT (blue) mice for strong connectivity. No effect on weak connectivity EC ratio was found. Dotted line represents no preference (ratio = 1). TBI = traumatic brain injury; CNO = clozapine n-oxide; fUS = functional ultrasound; ROI = region of interest; CBV = cerebral blood volume; S1Bf = primary somatosensory cortex, barrel field; S1Tr = primary somatosensory cortex, trunk region; RSV = ventral retrosplenial cortex; RSD = dorsal retrosplenial cortex; HIP = hippocampus; WD = weighted degree; EC = eigenvector centrality. See Supplementary Data S4 for statistical details. In **(b)** Atlas source: Paxinos, G. & PhD, K. B. J. F., MA. *The Mouse Brain in Stereotaxic Coordinates*. (Elsevier Science, 2007). Bar graphs represent Mean ± SEM. Each circle represents an individual sample.

Functional connectivity analysis of a selection of ROIs^42^ (Fig 7b,f) revealed that TBI reduces resting-state functional brain connectivity at P45 (Fig 7g,h). Intra-hemispheric connectivity of the primary somatosensory (S1) cortex regions (the direct “TBI impact area”) was reduced in TBI animals compared to sham (Fig 7j). This reduced connectivity in TBI was observed in the ipsilateral, but not the contralateral hemisphere (Fig 7j,k), reflecting a unilateral long-term impact of the TBI injury. Inter-hemispheric cortical connectivity was similarly reduced in TBI versus sham mice (Fig 7l). Correlation matrix analysis specifically revealed a loss of connectivity between S1TrL-RSDL, S1BFL-R, S1BFL-S1TrL, and RSDL-S1TrR, and a trend between RSVL-RSVR regions (Fig 7m; L refers to ipsilateral hemisphere, R to contralateral hemisphere. See figure legend for a full description of acronyms). Hippocampus connectivity was unaffected by TBI. Early-life oxytocin exposure after TBI was able to restore functional connectivity within the ipsilateral cortical regions (Fig 7j), and between hemispheric cortical regions (Fig 7l), compared to untreated TBI animals. This was specifically linked to increased connectivity between S1TrL-RSDL, RSDL-S1TrR and RSVL-R regions (Fig 7m). Similar effects of TBI and oxytocin were found on connectivity in corpus callosum regions (Fig S6). These data suggest that oxytocin release improves long-term functional connectivity of TBI-affected regions.

Next, we performed network analysis and visualization using Gephi^43^ on fUS connectivity between 212 ROIs across 3 coronal planes, to explore neural connectivity in more depth. From these neural networks, we calculated weighted degree measures of connectivity between structures, and eigenvector centrality measures of structure importance (or “centrality”) for the neural network. We focused network analysis on the isocortex structural hub, which included 56 ROIs across the 3 coronal planes recorded. The isocortex was divided into 5 substructures: sensorimotor cortex and retrosplenial cortex – identified as the direct TBI impact locations – and the peripheral auditory, perirhinal and association cortices. The isocortex neural network was filtered based on edge weights and analyzed separately for strong and weak connections. Fig 7n shows the average isocortex neural networks for Sham, TBI and TBI+OXT groups, divided into strong and weak connections, and color-coded for the 5 substructures. Node size represents eigenvector centrality, and edge size represents weighted degree. The networks are visualized using the Fruchterman-Reingold force-directed layout algorithm^44^, which pulls well-connected nodes to the center and pushes less-connected nodes to the perimeter.

Network analysis revealed that TBI mice had significantly fewer strong connections in the isocortex compared to sham, showing a reduction in both weighted degree of connections (Fig 7o), and eigenvector centrality measures of nodes (Fig 7p). These effects were found in every isocortex sub-structure, which suggests that TBI can alter connectivity of brain regions transcending the direct impact zone. Oxytocin treatment reversed the weighted degree of strong connections in the perirhinal cortex, but not in other substructures (Fig 7s). TBI+OXT mice showed a rescue of centrality in the isocortex as a whole (Fig 7p), as well as for the sensorimotor and retrosplenial substructures (Fig 7t). This suggests that endogenous oxytocin release creates a shift in which structures are a central part of the neural network after TBI.

Like strong connectivity, weak connections and anticorrelations are an important part of intrinsic brain connectivity^45^. The weighted degree of weak connections of the isocortex was increased in TBI versus sham, and rescued by oxytocin (Fig 7q). TBI mice showed an increase in weak connections in the retrosplenial, somatosensory and auditory networks (Fig 7u). The increased weak connectivity of these networks was reversed to sham-equivalent levels in TBI+OXT mice (Fig 7u). Treatment group did not affect centrality of weak isocortex connections for the whole network (Fig 7r), likely because weak connections lack power to affect the network as a whole. These data suggest that TBI mice increase their weak connections of the isocortex, at the cost of strong connections, and that this pattern is reversed by oxytocin treatment. No effects on connectivity for other structural hubs were found.

Given the unilateral damage of the TBI impact seen at P7, we wondered if there were differences in centrality between the ipsilateral and contralateral cortex for the 3 experimental groups. In a typical brain, both hemispheres contribute relatively equally to the neural network^46^. This was illustrated by the sham group, who demonstrated equal ratio of ipsilateral versus contralateral eigenvector centrality for isocortex strong connectivity (value 1 indicates equal balance; Fig 7v). TBI mice showed a reduction of this ratio, favoring the contralateral hemisphere in network centrality (Fig 7v). TBI+OXT mice had a fully restored equal balance in centrality between ipsilateral and contralateral isocortex regions, similar to sham mice (Fig 7v). This change in ipsilateral centrality by TBI and oxytocin was a spread effect, and not attributed to a specific substructure.

2.7 Oxytocin release rescues long-term behavioral deficits induced by TBI

Cognitive-behavioral problems are a well-established clinical outcome of pediatric TBI^6^. We subjected the 3 experimental groups to a behavioral test battery at the juvenile (P14) and adult age (P45) (Fig 8a). Morphometric features of animals were comparable among experimental groups as neither TBI nor oxytocin release affected body growth compared to sham (Fig 8b). The homing test (Fig 8c) is used as an early-life indicator of sociability and attachment, as it applies to the pup’s ability to recognize maternal scent^47^. Unlike sham mice, TBI mice did not show a preference for the home-cage nest bedding compared to bedding from an unfamiliar nest at P14 (Fig 8d). This impaired early social behavior persisted in the adult age, with TBI mice showing a reduced preference for the social stimulus during the 3-chamber test at P45, compared to sham (Fig 8h,i). TBI mice further demonstrated hyperactive behavior during the open field test at P45, showing increased average velocity and total distance traveled compared to sham (Fig 8e-g). No effect on time spent in the center was found.

**Fig. 8:**
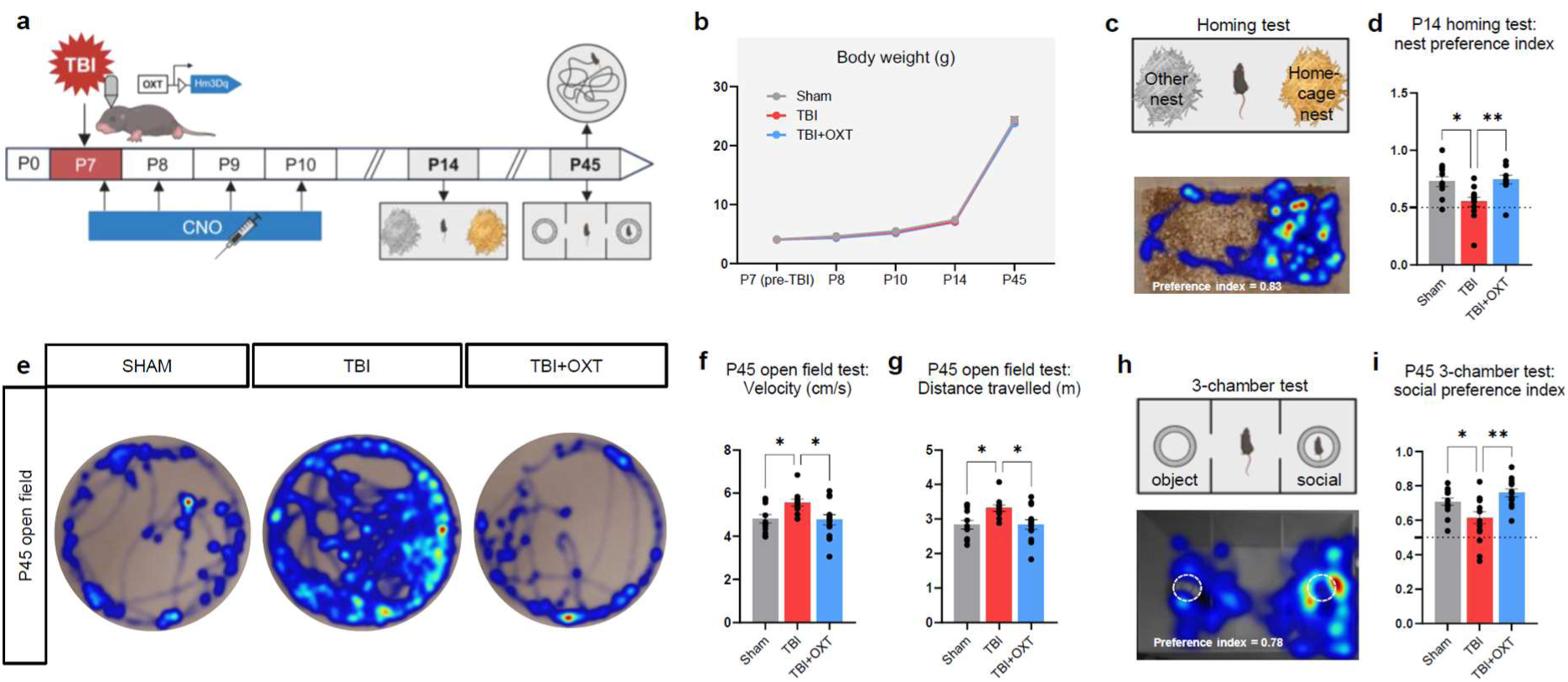
Long-term behavioral deficits of TBI are rescued by oxytocin treatment. **a)** Graphical timeline of behavioral experiments after P7 TBI and P7-P10 oxytocin treatment via CNO injections. **b)** Body weight was not affected by TBI or oxytocin treatment. **c)** Graphical representation and representative sham-level performance of the P14 homing test. **d)** Quantification of the P14 homing test nest preference index (PI) showed a decreased nest preference in TBI mice and a rescue in TBI+OXT mice. Dotted line represents no nest preference (PI = 0.5). **e)** Representative behavior of sham (left), TBI (middle) and TBI+OXT (right) mice during the P45 open field test. **f,g)** Quantification of the P45 open field test revealed an increase in velocity **(f)** and distance traveled **(g)** in the TBI mice, and a rescue of both behaviors in the TBI+OXT treatment group. **h)** Graphical representation and representative sham-level performance of the 3-chamber test at P45. **i)** Quantification of the P45 3-chamber test showed a decrease in social preference in the TBI mice and a rescue in the TBI+OXT mice. Dotted line represents no social preference (PI = 0.5). In **(b)** Mixed-effects analysis (F(8,161) = 0.34, *p* = .95, Sham: *n* = 15, TBI: *n* = 16, TBI+OXT: *n* = 14). In **(d)** ANOVA (F(2,34) = 6.91, \*\**p* = .003, Sham: *n* = 13, TBI: *n* = 13, TBI+OXT: *n* = 11). Tukey’s (Sham vs. TBI: \**p* = .01; Sham vs. TBI+OXT: *p* = .95; TBI vs. TBI+OXT: \*\**p* = .0067). In **(n)** ANOVA (F(2,38) = 0.34, *p* = .71, Sham: *n* = 13, TBI: *n* = 15, TBI+OXT: *n* = 13). In **(f)** ANOVA (F(2,33) = 4.54, \**p* = .0181, Sham: *n* = 12, TBI: *n* = 11, TBI+OXT: *n* = 13). Tukey’s (Sham vs. TBI: \**p* = .043; Sham vs. TBI+OXT: *p* = .98; TBI vs. TBI+OXT: \**p* = .026). In **(g)** ANOVA (F(2,33) = 4.73, \**p* = .016, Sham: *n* = 12, TBI: *n* = 11, TBI+OXT: *n* = 13). Tukey’s (Sham vs. TBI: \**p* = .029; Sham vs. TBI+OXT: *p* = .99; TBI vs. TBI+OXT: \**p* = .029). In **(i)** ANOVA (F(2,39) = 7.21, \*\**p* = .0022, Sham: *n* = 14, TBI: *n* = 15, TBI+OXT: *n* = 13). Tukey’s (Sham vs. TBI: \**p* = .049; Sham vs. TBI+OXT: *p* = .405; TBI vs. TBI+OXT: \*\**p* = .0018). Bar and line graphs represent Mean ± SEM. Each circle represents an individual sample.

Early oxytocin release reversed the long-term impairments in social behavior induced by TBI. TBI+OXT mice showed a fully restored preference for their home-cage nest bedding at P14 (Fig 8d), and for the social stimulus at P45 (Fig 8i). The hyperactivity of TBI mice found at P45 was reversed to sham-like levels in TBI+OXT mice (Fig 8f,g). These data suggest that acute oxytocin treatment following injury has long-lasting positive effects on behavioral issues caused by pediatric TBI.

Following the long-term effects of early-life TBI on P45 brain functioning, we investigated the neuroinflammatory profile at P45. We examined long-term neuroinflammation with microglia and astrocyte immunohistochemistry, and with *in-vivo* PET-imaging and *ex-*vivo gamma counting of the 18kDa translocator protein TSPO (Figure S6) – a protein expressed by inflammatory glia^48^. TBI mice showed no increase in cortical and white matter microglia density (Fig S7c,d) and astrocyte signal density (Fig S7e,f), and cortical TSPO expression (Fig S7g-h) at P45 compared to sham, nor did oxytocin treatment have an effect. This suggests that the P45 neurobehavioral deficits found in our pediatric TBI model were not associated with chronic neuroinflammation, and could be primarily long-term consequences of TBI-induced acute neuroinflammation.

## 3. Discussion

This study shows that chemogenetic activation of oxytocinergic neurons dampens the neuroinflammatory response 24 hours post-injury in a model of pediatric TBI. Early oxytocin release mitigated immune and pro-inflammatory gene pathways in astrocytes, and improved cerebrovascular repair and brain development mechanisms in microglial cells. The modulation of acute phase neuroinflammation prevented long-term damage to subcortical and cortical white matter tracts, to resting-state functional connectivity loss in the isocortex, and ultimately to social behavior and hyperactivity disorders induced by TBI.

Pediatric TBI can cause long-term impairments to multiple aspects of neurocognitive and behavioral functioning. First, white matter injury is a prominent feature of clinical pediatric TBI, that is similarly well documented in preclinical research^42,49,50^. Consistent with previous studies^49,51^, we report widespread white matter injury, with reduced long-term axonal integrity, myelin integrity and fiber tract thickness in our TBI model. Oxytocin release spared corpus callosum thickness and axonal integrity in the external capsule. It also improved myelination in the cortical TBI impact region. In agreement, clinical studies demonstrate improved fractional anisotropy in the whole brain and specifically in the external capsule in preterm infants exposed to music therapy^21^ – known to elevate oxytocin release^19,20^. Second, alterations in the resting-state functional brain connectivity after pediatric TBI have been demonstrated in clinical and preclinical research^42,52–54^. Functional ultrasound (fUS) imaging has improved spatial and temporal resolution over conventional imaging techniques such as fMRI^41^, and can be used both in rodents and young infants^14,42,55^. fUS imaging showed improved long-term functional connectivity in the TBI isocortex after oxytocin release, in both ipsilateral intra- and inter-hemispheric connectivity, 35 days after the treatment was given. Similar improvements were demonstrated when Carbetocin – a long-acting oxytocin receptor (OXTR) agonist – was used in a model of perinatal brain injury in rats^14^. With network analysis, we revealed that oxytocin can shift the centrality of cortical hubs within brain networks after TBI, restoring equal balance between hemispheres. Hub centrality within the neural network is important for executive functioning and information processing^56^, which can be distorted in children with TBI^52^. Oxytocin mostly affected isocortex connectivity, which agrees with studies showing increased functional connectivity and network centrality in the primary somatosensory and retrosplenial granular cortex in adult mice after environmental enrichment stimulation^57^ – which is associated with increased oxytocin levels^58^. Third, pediatric TBI can cause long-term behavioral issues, including antisocial behavior^59^. In line with the pro-social effects of oxytocin in autism spectrum disorder models^60,61^, we found that oxytocin restored typical social behavior in TBI-injured animals. A previous study performing TBI at P11 in male and female rats similarly observed a rescue of adolescent sociability after intranasal oxytocin administration 1h before testing^62^, and a study of iPSC-NPC transplantation after TBI at P14 in male rats reports improved social behavior associated with increased levels of oxytocin and oxytocin receptor expression^63^. Consistent with previous preclinical studies^26,51,64,65^, we found that pediatric TBI causes hyperactive disorders at adulthood. This was prevented by oxytocin release, similar to previous reports^14^, which shows that the behavioral effects of oxytocin transcend the social domain.

The three P45 readouts assessed in this study paint a potential cohesive picture of the long-term effects of oxytocin following pediatric TBI. As a major component of neural communication, white matter damage is a predictor of poor behavioral performance in clinical pediatric TBI^66,67^, and has established links with functional connectivity alterations^53^. Our findings of improved integrity of corpus callosum tracts following oxytocin treatment evidence inter-hemispheric functional connectivity of cortical regions. Similarly, increased white matter integrity in the ipsilateral cortex is associated with improved functional connectivity in that region. Moreover, the blood tracing modality of fUS imaging revealed restored long-term cerebral blood flow in oxytocin-treated TBI mice. Vascular growth, or ‘angiogenesis’, improves recovery of brain function and is a known therapeutic target following TBI^68,69^. Improved vascular integrity likely contributes to improved blood variation correlation that determines the functional connectivity modality of fUS^41^. Changes in functional connectivity following pediatric TBI have shown to correlate with impaired behavioral and cognitive functioning in children, and can persist up to years after the injury^52,70^. The isocortex – specifically its prefrontal modules – contains the neural correlates underlying social behavior in mice^71^, and has positive correlations with pro-social effects of oxytocin^62^. Improved functional connectivity of the ipsilateral isocortex following oxytocin treatment could underly the socio-behavioral improvements overserved here.

We found that long-term neuro-behavioral improvements induced by oxytocin release just after TBI link to specific effects on neuroinflammation during the acute post-injury phase. The acute neuroinflammatory response is a major contributor to disease progression – not only in TBI but after any insult to the developing brain^9^. Oxytocin reduced apoptotic cell death and the density and amoeboid morphology of microglial cells 24h post-injury. Such anti-inflammatory effects are known to improve disease progression after brain insult^72^. Anti-inflammatory effects on microglia reactivity have been demonstrated before for oxytocin^14^, and the current study extends these effects to acute injury in the developing brain. In addition to the direct effects on the cortical impact region, oxytocin reduced acute microgliosis in the ipsilateral corpus callosum. Microgliosis in developing white matter can impair oligodendrocyte viability and development, which causes long-term myelination deficits^73^. This is associated with increased expression of toxic molecules including reactive oxygen species (ROS), and cytokines TNF-α and IFN-γ^74^, as well as loss of insulin-like growth factor 1 (Igf1) expression by white matter microglia – which stimulates growth and proliferation of oligodendrocyte precursors. Impaired oligodendrocyte development could explain the acute local microgliosis and long-term integrity loss found in white matter of TBI mice, a notion supported by decreased Wnt signaling we found in TBI microglia, which causes hypomyelination^34^. Previous studies report a rescue of oligodendrocyte maturation following OXTR agonism in a perinatal brain injury model in rats^14^. A similar mechanism could be of effect in the current study, where modulation of acute neuroinflammation affects neurobehavioral outcomes of TBI through myelination.

Our results describe a potential spatial effect of the oxytocin release, in which microglia located in white matter areas were less affected than cortical microglia 24h post-injury. This heterogeneity mirrored the more moderate therapeutic effects of oxytocin on long-term subcortical white matter microstructure, compared to the cortex. One possible explanation concerns cell density. Unlike adult mice, the P7 mouse brain has a high density of microglia in the corpus callosum, creating a “fountain of microglia” migrating from the ventricular zone^75^. As shown *in vitro*, a higher cell density reduces the phenotypic responsiveness of macrophage-like cells to environmental stimulation^76^. Another possibility is that white matter microglia have less innate responsiveness to oxytocin. Single cell and spatial transcriptomic studies have revealed the vast heterogeneity of microglia, and suggest that white and gray matter microglia have distinct transcriptomic profiles^77,78^. We found a notably higher degree of oxytocin receptor mRNA expression in cortical microglia, which could suggest that cortical microglia respond better to oxytocin treatment. This difference is potentially associated with the unequal distribution of axonal projections of hypothalamic oxytocin neurons in the brain, showing higher innervation of gray matter rather than white matter regions^79^. Such a spatial heterogeneity in microglial oxytocin responsiveness will be important to explore in future studies.

Transcriptomic profiling of microglia and astrocytes after TBI provides insights into the ongoing inflammatory processes. On a transcriptomic level, oxytocin positively regulated genes encoding for tissue repair and brain development processes in TBI microglia. It increased the expression of genes related to vascular recovery, which likely contributes to the long-term vascular improvements we found at P45. Moreover, oxytocin reversed the transcriptomic increase of epigenetic processes in TBI microglia. Following an inflammatory insult as associated with TBI, microglia can undergo epigenetic reprogramming that heightens their response to secondary stressors^80^. In humans, such a two-hit model of inflammation and stress can lead to increased anxiety-like and depressive behaviors in adulthood^81^. Our findings suggest that oxytocin potentially reduces this pre-conditioning, which could protect the brain against future secondary insults. In astrocytes, oxytocin exposure increased expression of synaptic pathways, which are involved in the development of neural connectivity^82^. More notable was oxytocin’s reduction of the inflammation and immune pathways upregulated after TBI. This was linked to type 1 interferon signaling, a pathway whose downregulation is a known neuroprotective following TBI^25^.

The transcriptomic signatures suggest a differential effect of oxytocin release on astrocytes and microglia in the acute phase after injury. First, the number of affected genes was 5 times higher in microglia than in astrocytes. Second, the transcriptomic changes induced by oxytocin showed a different profile in astrocytes and microglia. During neuroinflammation, astrocytes and microglia shift between pro- and anti-inflammatory activity, where anti-inflammatory functions are traditionally associated with tissue-repair and take over later in the post-injury timeline^83^. We found that oxytocin mostly reduced pro-inflammatory processes in astrocytes, and mostly increased anti-inflammatory processes in microglia. This double-edged and cell-specific effect suggests that oxytocin accelerates the microglial shift towards a neuroprotective phenotype. Both effects of oxytocin likely contribute to improved outcomes following TBI, however the effects on microglia show a more direct link to the improved long-term outcomes of brain functioning at adult age. Other anti-inflammatory therapies likewise describe a different effect on microglia and astrocytes during inflammation^84^, and increased anti-inflammatory rather than a suppressed pro-inflammatory effect on microglia post-TBI has been reported^85^. Glial crosstalk during neuroinflammation is a known complex mechanism^86,87^, further complicated by the fact that the dynamic post-TBI transcriptomic profiles of microglia and astrocytes are highly susceptible to injury severity^25^, and timeline^88^. A practical distinction in neuroinflammation therapy research is whether to decrease toxic pro-inflammatory activity or increase protective anti-inflammatory activity, a dual mechanism that has been studied to great extent *in-vitro*^89–91^. The current study reveals the potential of oxytocin combining the two mechanisms, depending on glia type.

Oxytocin as a treatment of neuroinflammation is gaining scientific interest^13–15^. Here, we present new findings that oxytocin has acute anti-inflammatory effects after trauma in the developing brain. Previous studies using other therapies have likewise demonstrated beneficial effects decreasing microgliosis and inflammation post-TBI. However, they focus on short-term effects^26,27,50,92,93^, or require many treatment sessions^65^ (including pre-treatment^88,94^). Our data shows long-term benefits that persist long after oxytocin modulation is discontinued, with only a few treatment sessions required. As neuroinflammation is a key symptom of most types of perinatal brain injury^95^, the current study may act as a proof-of-concept for other brain injuries observed in young children. Translation of preclinical treatments is a general challenge for the pediatric population, which is aided by suboptimal study methodologies^11^. Methodological characteristics of the current study could contribute to improve the translational potential of the therapeutic findings. Chemogenetic manipulation of endogenous oxytocin via the DREADD construct – rather than exogenous oxytocin administration – mimics the naturally elevated oxytocin levels observed in clinical studies of music therapy, skin-to-skin contact, and environmental enrichment in infants^17–19^. Similarly, the closed-head weight-impact model of TBI is more appropriate to study pediatric TBI than the often used penetrating controlled-cortical-impact models of TBI^10^, given that the nature of TBI in newborns and children is predominantly non-penetrating physical impact^2^. Finally, clinical oxytocin elevation therapies are non-invasive and do not compete with traditional medication^17,20^. It thus has the potential to be implemented in addition to other clinical treatments. It should be noted that this study only included male mice – due to the increased occurrence and susceptibility of pediatric TBI seen in boys^96^. Given the increasing evidence for sexual dimorphism in early-life microglia^77,97^, in the inflammatory response to and outcomes of TBI^98^, and in the trajectory of the developing oxytocin system^99^, consideration of findings from this study should be limited to males and invite future assessment in females.

In conclusion, this preclinical study describes a biological foundation for a protective effect of endogenous oxytocin activity on brain development following experimental pediatric TBI. We show a multi-faceted improvement on brain functioning that persisted long after the treatment period, linked to changes to the microglial and astrocytic profile in the acute phase after injury. This identifies oxytocin as a novel, easy-to-implement treatment option for pediatric TBI in the clinical practice, which invites further exploration into its use.

## 4. Methods

### 1 Animals

Cre-dependent Hm3Dq-DREADD (Designer Receptors Exclusively Activated by Designer Drugs) mice (B6N;129-Tg(CAG-CHRM3*,-mCitrine)1Ute/J^28^, Jackson stock #026220) were crossed with Oxytocin-Ires-Cre mice (B6;129S-Oxttm1.1(cre)Dolsn/J, Jackson stock #024234) to create Oxt-Hm3Dq mice (B6;129S-Oxt-Cre;R26-LSL-hM3Dq-DREADD), which express the excitatory DREADD under the oxytocin promotor gene. This genetic profile allowed temporally increased neuronal firing of endogenous oxytocin neurons following injections of clozapine-N oxide solution (Enzo Life, BML-NS-105-0025).

Mice were housed in the SPF area in IVC-filtered top cages, under a light-dark cycle of 12 hours, with water and food access *ad libitum*. Room temperature and hygrometry were between 20-24°C and between 30% and 70%, respectively. Experimental cages included 2 dams with pups, and pups were randomly assigned to experimental groups. To prevent litter effects, each litter was divided into at least 2 experimental groups and every experiment consisted of at least 3 litters of animals. All surgeries and experiments were performed at the same hour across different litters. Only male mice were used in this study, due to the increased occurrence and susceptibility of pediatric TBI seen in male infants in the clinic ^96^.

This study was approved by Ethical Committee for Animal Experiments of the University of Geneva and the Veterinary Office of Geneva (cantonal no. GE155A; national no. 34289). The animal facility of the UNIGE Faculty of Medicine is approved by the General Direction on Health of Geneva and meets all the requirements defined by Swiss regulations and laws.

### 2 Pediatric traumatic brain injury model

A model of closed-head weight-drop impact acceleration injury to male P7 mice was used, adapted from Jacquens et al., 2024^42^. In brief, P7 mice were anesthetized with 4% isoflurane, confirmed by the loss of the toe-pinch reflex. An incision was made to the scalp to expose the skull, and the head was placed under a trauma contusion device consisting of a stainless-steel cylinder, which guides a 10 g weight falling onto a foot plate (2 mm diameter; RWD Stereotaxic Model 68093). The foot plate was positioned above the TBI location on the brain: on the left parietal bone, 1mm lateral and 2 mm anterior to lambda. The footplate was lowered until it contacted the skull, and then it was further depressed by 0.5 cm. Once fixated, the 10 g weight was dropped from a 12 cm distance onto the foot plate to induce the weight-drop impact acceleration injury to the skull. After the procedure, the scalp was closed with surgical glue (3M Vetbond 1469SB). Pups were kept under an infrared lamp until they woke up, after which they were put back in their home-cage. Before reintroduction in the home-cage, pups were gently rubbed with home-cage bedding to ensure successful re-acceptance by the dam. Sham-operated mice were anesthetized, their skull surface was exposed and then their skin was closed with surgical glue. The TBI injury was administered by the same experimenter to assure consistency across samples.

### 3 Pharmacological increase of endogenous oxytocin

Mice of the TBI-oxytocin treatment group received Clozapine-N oxide (CNO) injections to activate the excitatory DREADDs on oxytocin neurons, which causes a temporary increase in endogenous neuronal oxytocin firing^100^. CNO was dissolved in NaCl at a concentration of 10mg/kg and injected intraperitoneally (i.p.) at a volume of 10 μl. Mice from the oxytocin treatment group received daily CNO injections starting at P7, 4 hours post-TBI (when major inflammatory cytokines are peaking^101^), and subsequently every morning at P8, P9 and P10. Non-treated mice received NaCl injections at the same time points. CNO solutions were prepared fresh daily.

This study had 3 experimental groups: Sham-operated mice, receiving NaCl injections (“Sham mice”), TBI-operated mice, receiving NaCl injections (“TBI mice”), and TBI-operated mice, receiving oxytocin-increasing CNO injections (“TBI+OXT mice”).

### 4 Evans Blue assay of cerebrovascular rupture

To assess blood-brain-barrier integrity loss in the TBI model, and to quantify rupture of cortical vasculature in response to the TBI procedure, we performed an Evans Blue assay at P7. Mice were i.p. injected with 50 μl of 1% Evans Blue (Sigma-Aldrich E2129) and put back in the home-cage to let the solution circulate and reach the brain vasculature. 4 hours post-injection, mice were subjected to TBI or Sham-operation. Mice were sacrificed 4 hours post-surgery with decapitation after pentobarbital anesthesia (150 mg/kg). Brains were immersed in 4% PFA for 48 hours, washed with PBS1x and imaged using iBright FL1500 Imaging System (ThermoFisher) at AlexaFluor 633 wavelength. Cerebrovascular rupture was assessed with ImageJ as an invert measure of ipsilateral hemisphere Evans Blue immunodensity fold change for TBI mice versus Sham littermate controls.

### 5 Immunohistology

Immunohistology was performed at P8 (24h post-TBI) and P45 (38 days post-TBI). Animals were anesthetized with i.p. injections of pentobarbital (150 mg/kg), confirmed by the loss of the toe-pinch reflex, and transcardially perfused with PBS and 4% paraformaldehyde (PFA) in PBS1x. Brains were post-fixated overnight in 4% PFA, and embedded in 15% sucrose (morning) and 30% sucrose (afternoon) solution the following day. Brains were snap-frozen in isopentane at -45°C and stored at - 20°C until cutting. Brains were cut with a cryostat into free-floating coronal sections of 50 μm and stored in PBS+0.3%Azide solution. Coronal sections corresponding to slide 70 to 75 of Allen brain atlas (atlas.brain-map.org) were selected for immunofluorescent assays, corresponding to the Bregma level of TBI impact.

#### 5.1 Immunofluorescence

Free-floating brain sections were blocked with 0.2% Triton+10% BSA in PBS1x for 1 hour. Following a 5 min wash in PBS, sections were incubated overnight at 4°C with primary antibodies in a solution of 0.2%Triton+3% BSA in PBS1x. Primary antibodies included rabbit anti-cleaved-caspase 3 (1:200; Cell Signaling D175), goat anti-Iba1 (1:500; Abcam ab5076), rabbit anti-Iba1 (1:1000; Fujifilm Wako 019-19741) and rat anti-CD68 (1:1000, Bio-rad FA-11), to assess (neuronal) cell death, microglia reactivity and their phagocytic activity at P8, and mouse anti-MBP (1:500, Sigma-Aldrich MAB382) was used to investigate myelination at P45. Following 3x 10 min washes in PBS1x, sections were incubated with secondary antibodies in a solution of 0.2%Triton+10% BSA in PBS1x for 1 hour and 30 minutes. Secondary antibodies were goat anti rabbit 594 (1:500, Abcam ab150080), goat anti rat 488 (1:500, Abcam ab150165), goat anti mouse 647 (1:500, Abcam ab150115), donkey anti rabbit 488 (1:500; Abcam ab150061), chicken anti goat 647 (1:500, Invitrogen A21469). After a final wash in PBS1x, sections were mounted and coverslipped on microscopy slides (ThermoFisher) with flouromount medium that includes DAPI (Abcam ab104139). Slides were stored at 4°C.

#### 5.2 Image acquisition

For measuring the degree of apoptotic cell death and microglia response at P8, brain sections were imaged with a ZEISS Axioscan.Z1 (Zeiss, Oberkochen, Germany) at 10x. Other immunohistochemistry assays were imaged using a ZEISS LSM 800 Airyscan confocal microscope (Zeiss, Oberkochen, Germany), at different magnitudes. All images were taken in the hemisphere ipsilateral to the brain injury. Micrographs were acquired by researchers blinded to experimental conditions.

For measuring the number of microglia and CD68 signal immunodensity, 3×4 tiles of 10x magnification were taken in two ROIs: the medial somatosensory cortex (layers I-VI), and the medial corpus callosum including the cingulate bundle. Both ROIs were taken at a horizontal localization superimposing the hippocampal CA1 region.

For single-cell microglia morphological analysis, 4 40X z-stack images (0.23 μm step size) were taken per animal, in 2 ROIs: in the medial somatosensory cortex layers IV/V, and at the medial corpus callosum including the cingulate bundle (superimposing the hippocampal CA1 region).

For analysis of corpus callosum thickness, 3×3 10x tiles were taken in the center of the structure, at the hemispheric division. For analysis of the external capsule, 3×4 10x tiles were taken. For analysis of cortical myelination, 3 40x z-stack images (0.23 μm step size) were taken per animal, in adjacent regions of cortical layers IV/V superimposing the hippocampal CA1 and CA2 regions.

#### 5.3 Image analysis

Images were analyzed using FIJI/ImageJ^102^ and QuPath^103^. Quantifications were performed by researchers blinded to experimental conditions.

##### 5.3.1 Apoptotic cell death and microglial proliferation

To quantify the degree of apoptotic cell death in the S1 cortex and white matter (WM), we counted the number of cleaved caspase-3^+^ (c-casp3) cells, and the proportion of NeuN^+^/c-casp3^+^ cells. We also counted the number of microglia that were touching or engulfing c-casp3^+^ cells. Microglia density in the S1 and WM was quantified as the number of Iba1^+^ cells in the ROIs. Data from 3 micrographs per animal were averaged.

##### 5.3.2 Morphological analysis of single microglia

To assess the morphological characteristics of microglia on a single-cell level, individual microglia were isolated and reconstructed from the 40x z-stack micrographs, and saved as binary images for further analysis with the Shape Filter plugin^104^ and Sholl plugin^105^ of ImageJ.

###### 5.3.2.1 Shape filter analysis

Shape filter analysis ^104^ was performed on individual microglia to extract detailed parameters of overall cell morphology: Cell perimeter (CP) in μm, Cell density (CD), which is the ratio between cell area and cell perimeter, Cell convexity, which is the perimeter of the convex hull divided by the perimeter of the cell. Finally, a “Cell Circularity” measure was calculated with the following formula: circularity = 4π x CA / CP^2^. Measures from 4-5 microglia were averaged per ROI per animal.

###### 5.3.2.2 Sholl analysis

To measure the ramification of microglial cells, Sholl analysis^105^ was performed on individual microglia. Sholl analysis was performed with a 2 μm radius step size, to generate a readout of number of intersections per radius step. 4-5 microglia were averaged per ROI per animal.

##### 5.3.3 Microglia phagocytic compartment

Phagocytic engulfment of dead or dying cells by microglia was assessed as the number of c-casp3^+^-Iba1^+^ touching cells, determined by manual counting. To further assess the phagocytic activity, the CD68^+^ microglial compartment was measured as fluorescence density from a 3×4 10x tile micrograph. For each animal, an integrated CD68 density measure was calculated from a selected ROI. Fluorescence of 3 separate background readings per ROI was averaged and subtracted from the CD68 integrated density while controlling for ROI size using the formula: Corrected CD68 fluorescence = Integrated Density – (Area size of selected region x Mean fluorescence of background readings). Corrected CD68^+^ fluorescence was normalized on the Sham group and reported as a fold change.

##### 5.3.4 White matter microstructure

Analysis of white matter microstructure at P45 was performed in ImageJ and based on the methods reported previously^40^. Subcortical white matter and the integrity of myelinated axons in the cortex were based on myelin base protein (MBP) immunoreactivity.

###### 5.3.4.1 Subcortical white matter

Subcortical white matter microstructure was assessed as the size (μm) of the medial corpus callosum and the external capsule based on 10x micrographs.

###### 5.3.4.2.1 Cortical MBP immunodensity and MBP^+^ area

The MBP immunodensity and MBP^+^ area measures were based on 40x z-stack micrographs taken as 3 adjacent regions in the somatosensory cortex (cortical layers IV/V). MBP immunodensity was measured as the mean signal intensity measure of the whole micrograph, subtracted by the background signal intensity. The MBP immunodensity was normalized on Sham group values, which were set on 100%. For each animal, measures from the 3 micrographs were averaged.

For the MBP^+^ area analysis, a threshold for the MBP signal was applied to the whole micrographs, creating a binary image of all MBP^+^ tissue and removing the background. The area proportion of MBP^+^ signal was then calculated. Measures from 3 micrographs were averaged per animal.

###### 5.3.4.2.2 Cortical MBP fiber coherency

MBP fiber coherency is an inverted measure of fiber complexity ^40^. This was assessed in 3 adjacent 40x z-stack micrographs of the ipsilateral cortex, with the ‘OrientationJ measure’ function of the ImageJ plugin OrientationJ^106^, as described previously^40^. Data from 3 micrographs were averaged per animal.

### 6 Transcriptomic profiling of glia

#### 6.1 Magnetic isolation of microglia and astrocytes

To enable *in-vivo* assessment of the acute microglia and astrocyte transcriptome following TBI and the oxytocin treatment, microglia and astrocytes were sorted from P8 mice using magnetic cell sorting technology as described previously^107^. Briefly, P8 pups were anesthetized with pentobarbital i.p. injection (150 mg/kg) and transcardially perfused with PBS to remove macrophages. Brains were dissected, olfactory bulb and cerebellum removed, and the hemisphere ipsilateral to the injury was selected for further processing. Brain tissue was dissociated using the MACS® neural dissociation kit (Miltenyi Biotech). Microglia cells were isolated from brain homogenates via incubation with magnetic anti-mouse CD11b microbeads (Miltenyi Biotech) and collected using the MultiMACS® cell separator apparatus (Miltenyi Biotech). A separate sorting experiment was executed to collect astrocytes (using anti-mouse ACSA-2 microbeads). Sorted microglia and astrocytes were stored at -80°C.

#### 6.2 Microglia real-time qPCR

For validation of the TBI model, RNA was extracted from P8 MACS-sorted microglia using the Nucleospin RNA Plus XS extraction kit (Macherey-Nagel). RNA quantity was measured with Nanodrop 2000 and was normalized across samples. RNA was subjected to reverse transcriptase using the MLV-RT kit (Promega; M1701). Real-time quantitative transcription polymerase chain reaction (RT-qPCR) was performed in duplicates with PowerUp™ SYBR™ Green Master Mix (ThermoFischer) for 40 cycles with a 2-step program (15 sec of denaturation at 95°C and 1 min of annealing/extension at 60°C). mRNA amplification values were normalized on Rps18 housekeeping gene and calculated as mRNA fold change measures normalized on the Sham group following the ΔΔCt analysis method. Gene expression of a selection of pro-inflammatory cytokines was assessed, based on previous findings^42^. Primers were designed with Primer-BLAST^108^. Primer sequences are provided in Table S2.

#### 6.3 Microglia and astrocyte RNA sequencing

RNA was extracted from P8 sorted microglia using the Nucleospin RNA Plus XS extraction kit (Macherey-Nagel). RNA quantification was performed with a Qubit fluorometer (ThermoFisher Scientific) and RNA integrity assessed with a Bioanalyzer (Agilent Technologies). The microglial RNA was sequenced in bulk with Illumina NovaSeq 6000 next-generation sequencer. Quality control was performed with FastQC (v.0.1.1.9), and the reads were mapped to the reference mouse genome (Ensembl / GRCm39) using STAR aligner (v.2.7.10b).

MACS cell sorting purity was assessed with Expression Weighted Cell-type Enrichment analysis^29^ op the top 250 expressed genes in the Sham group, based on normalized gene expression. The analysis was run with default parameters, with the number of repetitions for bootstrap enrichment set to 10.000. Enrichment significance was set at p < .05.

Differential gene expression (DEG) analysis was performed with edgeR (v.3.42.4). Lowly expressed genes (< 10 raw counts) were filtered out, and gene expression was normalized according to the library size and RNA composition. Differentially expressed genes were estimated using a quasi-likelihood (QL) F-test with rigorous type I error rate control. Corrections for multiple testing were performed with False Discovery Rate (FDR) and the Benjamini & Hochberg correction (BH). Significant genes were filtered on cut-off values of ≥ 1.5 fold-change and p < .05. DEG analysis was performed for the comparisons TBI versus Sham, and TBI+OXT versus TBI. Unidentified DEGs were excluded from the results.

Gene set enrichment analysis on differentially expressed genes was performed using DAVID Functional Annotation Tool (DAVID Knowledgebase v2024q1, National Institutes of Health). The upregulated DEGs were analyzed separately from the downregulated DEGs. Databases of biological processes and KEGG pathways were investigated. Pathways with a p value of < .05 were considered.

The same methodology was applied to astrocyte RNA sequencing, with the exception that a batch effect was discovered for astrocyte samples with PCA analysis, which was corrected for in downstream RNA Seq analyses.

The RNA Seq experiments were performed at the iGE3 Genomics Platform of the University of Geneva (https://ige3.genomics.unige.ch).

### 7 *Ex-vivo* brain structure assessment with MRI-diffusion tensor imaging

We performed microscopic diffusion analysis of white and grey matter in the ipsilateral hemisphere with *ex-vivo* MRI imaging to assess long-term microstructural brain development. At P45, animals were anesthetized with i.p. injections of pentobarbital (150 mg/kg), confirmed by the loss of the toe-pinch reflex, and transcardially perfused with PBS and 4% paraformaldehyde (PFA) in PBS1x. Brains were kept in PFA overnight, then changed and kept in PBS1x for 6 weeks until scanning.

#### 7.1 Image acquisition

*Ex-vivo* MRI experiments were performed with a 2.5 cm diameter birdcage coil, on an 9.4T/31cm actively shielded horizontal-bore magnet (Magnex Scientific, Yarnton, UK), B-GA12S HP shielded gradient set (114mm ID, 660 mT/m peak strength and 4570 T/m/s slew rate, Bruker BioSpin, Ettlingen, Germany) interfaced to a Bruker BioSpec console with AVANCE NEO electronics running ParaVision 360 v3.5. A multi-b-value shell protocol was acquired using a spin-echo sequence (FOV = 21 × 16 mm^2^, matrix size = 120 × 80, 20 slices of 0.6 mm, 4 averages with TE/TR = 22/2000ms). 82 DWI were acquired, 6 b0 images and 76 separated in 3 shells (noncollinear and uniformly distributed in each shell with δΔ = 4.5/12 ms) with number of directions/b-value in s/mm^2^: 16/1500, 30/3000 and 30/5000.

#### 7.2 Microscopic diffusion analysis with DTI and NODDI techniques

Diffusion data processing started by MP-PCA denoising^109^ then the diffusion tensor was estimated using a weighted linear least squares algorithm^110^. Diffusion weighted data were fitted using the AMICO model^111^. Several brain regions were assessed in the ipsilateral hemisphere, ROIs were delineated on the fractional anisotropy maps. We calculated DTI derived parameters, including Axial diffusivity (AD), representing axonal integrity, Radial diffusivity (RD), representing myelin sheath integrity, and Fractional anisotropy (FA), reflecting axonal diameter, fiber density and myelination. We also calculated neurite orientation dispersion and density imaging (NODDI) metrics, including Neurite density index (NDI), relating to the packing density of axons or dendrites, Orientation dispersion index (ODI), an inverse measure of fiber alignment, and Free water fraction (FWF), which reflects contamination of the CSF. These parameters were calculated for 5 Ipsilateral ROIs for each animal: corpus callosum (CC), cingulum (Cg), external capsule (EC), primary motor cortex (M1Cx) and primary somatosensory cortex (S1Cx). Data were averaged for the coronal planes and statistically tested between groups using one-way ANOVA tests with Tukey’s post-hoc.

### 8 *In-vivo* brain connectivity imaging with functional ultrasound (fUS)

We performed *in-vivo* resting-state functional ultrasound (fUS) imaging of the brain at P45 as a readout of long-term functional brain connectivity. This method uses cerebrovascular blood volume (CBV) variations to analyze the correlation of resting-state neuronal activity, between ROIs in the same (intra-hemispheric connectivity) and opposite hemispheres (inter-hemispheric connectivity)^41^. The mouse P56 coronal brain atlas from Allen Institute (atlas.brain-map.org) was used as ROI map.

#### 8.1 fUS data acquisition

The experiment was performed as previously described^14,42^. Briefly, P45 animals were anesthetized with 1:10 Ketamine mixed with 1:20 Domitor in NaCl, injected i.p. at a concentration of 100 ul/kg body weight. After ensuring anesthesia by the loss of the toe-pinch reflex, the scalp was cut to expose the skull. The animal was placed in a stereotaxic frame, on top of a heating pad kept at 37°C. Ultrasound coupling gel was applied to the skull and the ultrasonic probe (Vermon, Tours, France) was positioned directly above the parietal bone.

An initial 3D anatomic scan was performed for every mouse to function as a reference atlas to align the ROI map to. The probe was vertically positioned directly above Bregma level -1. From this position, a 10-minute ultrasonic acquisition scan was conducted at a frame-rate of 500 Hz, with tilted plane waves oriented at 5 different angles to allow optimal detection of blood volume variation.

#### 8.2 fUS data analysis

##### 8.2.1 Connectivity correlation analysis

For each animal, CBV patterns were transformed into connectivity correlation matrices between a selection of 10 ROIs of the atlas, based on previous findings^42^: barrel field of the somatosensory cortex (S1BF), trunk region of the somatosensory cortex (S1Tr), ventral retrosplenial cortex (RSV), dorsal retrosplenial cortex (RSD) and CA1 region of the hippocampus (HIP), in both hemispheres. We analyzed connectivity between these regions, and for connectivity in the cortex (averaging the S1Tr, S1BF, RSV, and RSD regions) in the ipsilateral and contralateral hemispheres. For statistical testing between groups, the Pearson correlation coefficients were transformed using a Fisher z transformation, and tested with One-way ANOVAs.

##### 8.2.2 Neural network analysis

To perform a holistic network analysis of brain connectivity, we measured CBV variations in 212 ROIs across 3 coronal planes (Bregma -1, -1.5, -2). This created a large connectivity matrix that was loaded into Gephi^43^ where it was transformed into a 3D neural network of nodes (ROIs) and edges (connections between ROIs). We categorized the 212 ROIs into cerebral structures (e.g., “sensorimotor cortex”) and transcending structural “hubs” (e.g., “isocortex”), to enable assessment of larger collections of structures in the network. See full list of ROIs and their categorization in Table S1.

Neural networks were filtered on structural hub, edge weight (strong *vs* weak connections), underlying structure and hemisphere. We analyzed for every structural hub and for a selection of underlying structures the total connectivity (as average weighted degree and average eigenvector centrality) and the ratio between ipsilateral and contralateral hemispheres for the eigenvector centrality measures. Weighted degree reflects the average strength of connections of the network, and eigenvector centrality also includes the strength of connecting nodes^112^ – meaning that it is a measure of how central the node is in the entire network. Statistical comparison between treatment groups was performed with One-way ANOVA with Tukey’s post-hoc assessment.

### 9 Behavior

Mouse behavior was tracked and analyzed using Ethovision^113^. Behavioral experiments were performed by the same researcher to ensure a consistent testing environment.

#### 9.1 Body growth

Body growth was measured at P7 (before TBI procedure), P8, P10, P14 and P45 to assess effects of the TBI and the oxytocin treatment on general well-being of the mouse.

#### 9.2 Homing test

At P14, the homing test was performed to analyze early-life sociability: a pup was placed in a 32×16×18 cm (lxwxh) cage with bedding and was allowed free exploration during 3 minutes. The cage was divided in length into 2 outer zones of 12 cm. The middle zone (8 cm) had fresh bedding, one zone on the far side had bedding from the pup’s home-cage nest, and the other far-side zone had bedding from an unfamiliar nest with pups. Time spent in each zone was calculated and converted to a preference index for the home-cage zone versus the unfamiliar nest zone. The cage orientation was alternated between trials and fresh bedding from the 2 nests was added between every trial to equalize odor strength between trials.

#### 9.3 Open field test

The open field test was performed at P45 to analyze general motor activity and anxiety. Animals were placed in a circular arena (35 cm diameter, 22 cm tall) and were allowed free exploration during 10 minutes. Readouts were velocity (m/s), total distance travelled (m), and time spent in the center zone (s) as an indirect measure of anxiety.

#### 9.4 Three chamber test

The three-chamber test was performed at P45 as an adult measure of sociability. In an arena divided in 3 connected chambers of 20×40×25 cm (lxwxh), the two outer chambers include either an empty mesh-bar enclosure (9 cm diameter, 17 cm tall) or an enclosure containing an unfamiliar mouse (one week younger than the experimental mouse). After a 10 min habituation phase, the 2 enclosures were placed and the animal was allowed free exploration during 5 minutes. Data was analyzed as time spent interacting with both enclosures (object versus social), by tracking the total time the animal’s nose was within 2 cm proximity from the enclosure. We analyzed the social preference score by calculating the time spent interacting with the social enclosure divided by the total time interacting with the social + empty enclosure. 10 Statistical analysis Statistical analysis was performed with Graphpad/Prism version 10.4.0 for Windows (GraphPad Software, Boston, Massachusetts USA, www.graphpad.com). All data was checked for outliers with the ROUT method (Q = 0.1%), and tested for normality using Kolmogorov-Smirnov test (α = .05) and QQ plot inspection. The *n* indicated in each figure legend represents sample size after outlier removal. Comparisons between 2 treatment groups were performed with Student’s t-test for normally distributed data and a Mann-Whitney U test for non-normally distributed data. Analyses of phagocytosis between cortex and white matter in the TBI group was performed with paired t-test. Analyses between all 3 treatment groups was performed with a One-way ANOVA with Tukey’s post-hoc for normal distributed data, or Kruskal-Wallis test with Dunn’s post-hoc for non-normally distributed data. Sholl analysis of microglia ramification was performed with a two-way ANOVA of treatment x sholl radius. Behavioral assays of repetitive assessments (e.g., body growth) were analyzed with a two-way ANOVA of treatment x postnatal age. The alpha was set at 0.05, and tested two-tailed. Unless specified otherwise, graphs show Mean + SEM values. Non-significant post-hoc comparisons are not shown in the graphs.

## 5. Data availability

All data necessary to evaluate the conclusions in this paper are included within the main text and Supplementary Materials. Source data are provided in a separate file.

## 7. Acknowledgemacents

This study was funded by the Swiss National Science Foundation (grant number 197462), and supported by La Fondation des Gueules Cassées and La Société Française d’Anesthésie et de Réanimation. Author ST was supported by the Swiss National Science Foundation (grant number 223414). We thank the CIBM Center for Biomedical Imaging for providing expertise and resources for the PET experiment. Figures were made with BioRender (https://www.biorender.com/), and SRplot^114^.

## Notes

### Competing Interest Statement

The authors have declared no competing interest.

## References

1. Dewan, M. C., Mummareddy, N., Wellons, J. C. & Bonfield, C. M. Epidemiology of Global Pediatric Traumatic Brain Injury: Qualitative Review. World Neurosurgery 91, 497–509.e1 (2016).

2. Araki, T., Yokota, H. & Morita, A. Pediatric Traumatic Brain Injury: Characteristic Features, Diagnosis, and Management. Neurol Med Chir (Tokyo) 57, 82–93 (2017).

3. Figaji, A. A. Anatomical and Physiological Differences between Children and Adults Relevant to Traumatic Brain Injury and the Implications for Clinical Assessment and Care. Front. Neurol. 8, (2017).

4. Simon, A. K., Hollander, G. A. & McMichael, A. Evolution of the immune system in humans from infancy to old age. Proc Biol Sci 282, 20143085 (2015).

5. Matcovitch-Natan, O. et al. Microglia development follows a stepwise program to regulate brain homeostasis. Science 353, aad8670 (2016).

6. Kooper, C. C. et al. Long-Term Neurodevelopmental Outcome of Children With Mild Traumatic Brain Injury. Pediatric Neurology 160, 18–25 (2024).

7. Kyösti, E. et al. Long-Term Quality of Life After Pediatric Traumatic Brain Injury Treated in the Intensive Care Unit. Pediatric Neurology 157, 50–56 (2024).

8. Figaji, A. An update on pediatric traumatic brain injury. Childs Nerv Syst 39, 3071–3081 (2023).

9. Jacquens, A. et al. Neuro-Inflammation Modulation and Post-Traumatic Brain Injury Lesions: From Bench to Bed-Side. International Journal of Molecular Sciences 23, 11193 (2022).

10. Kochanek, P. M., Wallisch, J. S., Bayır, H. & Clark, R. S. B. Pre-clinical models in pediatric traumatic brain injury-challenges and lessons learned. Childs Nerv Syst 33, 1693–1701 (2017).

11. Seyhan, A. A. Lost in translation: the valley of death across preclinical and clinical divide – identification of problems and overcoming obstacles. Translational Medicine Communications 4, 18 (2019).

12. Rokicki, J. et al. Oxytocin receptor expression patterns in the human brain across development. Neuropsychopharmacol. 47, 1550–1560 (2022).

13. Knoop, M. et al. The Role of Oxytocin in Abnormal Brain Development: Effect on Glial Cells and Neuroinflammation. Cells 11, 3899 (2022).

14. Mairesse, J. et al. Oxytocin receptor agonist reduces perinatal brain damage by targeting microglia. Glia 67, 345–359 (2019).

15. Yuan, L. et al. Oxytocin inhibits lipopolysaccharide-induced inflammation in microglial cells and attenuates microglial activation in lipopolysaccharide-treated mice. J Neuroinflammation 13, 77 (2016).

16. Sünnetçi, E., Solmaz, V. & Erbaş, O. Chronic Oxytocin treatment has long lasting therapeutic potential in a rat model of neonatal hypercapnic-hypoxia injury, through enhanced GABAergic signaling and by reducing hippocampal gliosis with its anti-inflammatory feature. Peptides 135, 170398 (2021).

17. Vittner, D. et al. Increase in Oxytocin From Skin-to-Skin Contact Enhances Development of Parent-Infant Relationship. Biol Res Nurs 20, 54–62 (2018).

18. Moberg, K. U., Handlin, L. & Petersson, M. Neuroendocrine mechanisms involved in the physiological effects caused by skin-to-skin contact – With a particular focus on the oxytocinergic system. Infant Behavior and Development 61, 101482 (2020).

19. Chanda, M. L. & Levitin, D. J. The neurochemistry of music. Trends Cogn Sci 17, 179–193 (2013).

20. Wulff, V. et al. The effects of a music and singing intervention during pregnancy on maternal well-being and mother–infant bonding: a randomised, controlled study. Arch Gynecol Obstet 303, 69–83 (2021).

21. Sa de Almeida, J., et al. Music enhances structural maturation of emotional processing neural pathways in very preterm infants. NeuroImage 207, 116391 (2020).

22. Konar, M. C., Islam, K., Sil, A., Nayek, K. & Barik, K. Effect of Music on Outcomes of Birth Asphyxia: A Randomized Controlled Trial. Journal of Tropical Pediatrics 67, fmab009 (2021).

23. Emery, L. et al. A randomised controlled trial of protocolised music therapy demonstrates developmental milestone acquisition in hospitalised infants. Acta Paediatrica 108, 828–834 (2019).

24. Ormston, K., Howard, R., Gallagher, K., Mitra, S. & Jaschke, A. The Role of Music Therapy with Infants with Perinatal Brain Injury. Brain Sci 12, 578 (2022).

25. Todd, B. P. et al. Traumatic brain injury results in unique microglial and astrocyte transcriptomes enriched for type I interferon response. Journal of Neuroinflammation 18, 151 (2021).

26. Semple, B. D. et al. Interleukin-1 Receptor in Seizure Susceptibility after Traumatic Injury to the Pediatric Brain. J Neurosci 37, 7864–7877 (2017).

27. Gu, X. et al. Pharmacologically Induced Hypothermia Attenuates Traumatic Brain Injury in Neonatal Rats. Exp Neurol 267, 135–142 (2015).

28. Zhu, H. et al. Cre dependent DREADD (Designer Receptors Exclusively Activated by Designer Drugs) mice. Genesis (New York, N.Y. : 2000) 54, 439 (2016).

29. Skene, N. G. & Grant, S. G. N. Identification of Vulnerable Cell Types in Major Brain Disorders Using Single Cell Transcriptomes and Expression Weighted Cell Type Enrichment. Front. Neurosci. 10, (2016).

30. Wang, L. et al. Glucose transporter 1 critically controls microglial activation through facilitating glycolysis. Mol Neurodegener 14, 2 (2019).

31. Xu, X. et al. HIF-1α participates in secondary brain injury through regulating neuroinflammation. Translational Neuroscience (2023) doi:10.1515/tnsci-2022-0272.

32. Jullienne, A. et al. Male and Female Mice Exhibit Divergent Responses of the Cortical Vasculature to Traumatic Brain Injury. J Neurotrauma 35, 1646–1658 (2018).

33. Chu, E., Mychasiuk, R., Hibbs, M. L. & Semple, B. D. Dysregulated phosphoinositide 3-kinase signaling in microglia: shaping chronic neuroinflammation. Journal of Neuroinflammation 18, 276 (2021).

34. Van Steenwinckel, J. et al. Decreased microglial Wnt/β-catenin signalling drives microglial pro-inflammatory activation in the developing brain. Brain 142, 3806–3833 (2019).

35. Jha, M. K., Jo, M., Kim, J.-H. & Suk, K. Microglia-Astrocyte Crosstalk: An Intimate Molecular Conversation. Neuroscientist 25, 227–240 (2019).

36. Guttenplan, K. A. et al. Knockout of reactive astrocyte activating factors slows disease progression in an ALS mouse model. Nat Commun 11, 3753 (2020).

37. Lee, H.-G., Lee, J.-H., Flausino, L. E. & Quintana, F. J. Neuroinflammation: An astrocyte perspective. Science Translational Medicine 15, eadi7828 (2023).

38. Colombo, E. & Farina, C. Astrocytes: Key Regulators of Neuroinflammation. Trends Immunol 37, 608–620 (2016).

39. Farizatto, K. L. G. & Baldwin, K. T. Astrocyte-synapse interactions during brain development. Current Opinion in Neurobiology 80, 102704 (2023).

40. van Tilborg, E. et al. A quantitative method for microstructural analysis of myelinated axons in the injured rodent brain. Sci Rep 7, 16492 (2017).

41. Macé, E. et al. Functional ultrasound imaging of the brain. Nat Methods 8, 662–664 (2011).

42. Jacquens, A. et al. Deleterious effect of sustained neuroinflammation in pediatric traumatic brain injury. Brain, Behavior, and Immunity 120, 99–116 (2024).

43. Bastian, M., Heymann, S. & Jacomy, M. Gephi: An Open Source Software for Exploring and Manipulating Networks. ICWSM 3, 361–362 (2009).

44. Fruchterman, T. M. J. & Reingold, E. M. Graph drawing by force-directed placement. Software: Practice and Experience 21, 1129–1164 (1991).

45. Fox, M. D. et al. The human brain is intrinsically organized into dynamic, anticorrelated functional networks. Proceedings of the National Academy of Sciences 102, 9673–9678 (2005).

46. Martínez, J. H., Buldú, J. M., Papo, D., Fallani, F. D. V. & Chavez, M. Role of inter-hemispheric connections in functional brain networks. Sci Rep 8, 10246 (2018).

47. Scattoni, M. L., Puopolo, M., Calamandrei, G. & Ricceri, L. Basal forebrain cholinergic lesions in 7-day-old rats alter ultrasound vocalisations and homing behaviour. Behavioural Brain Research 161, 169–172 (2005).

48. Tournier, B. B. et al. Fluorescence-activated cell sorting to reveal the cell origin of radioligand binding. J Cereb Blood Flow Metab 40, 1242–1255 (2020).

49. Dikranian, K. et al. Mild traumatic brain injury to the infant mouse causes robust white matter axonal degeneration which precedes apoptotic death of cortical and thalamic neurons. Experimental Neurology 211, 551–560 (2008).

50. Hanlon, L. A., Huh, J. W. & Raghupathi, R. Minocycline Transiently Reduces Microglia/Macrophage Activation but Exacerbates Cognitive Deficits Following Repetitive Traumatic Brain Injury in the Neonatal Rat. J Neuropathol Exp Neurol 75, 214–226 (2016).

51. Obenaus, A. et al. A single mild juvenile TBI in male mice leads to regional brain tissue abnormalities at 12 months of age that correlate with cognitive impairment at the middle age. Acta Neuropathol Commun 11, 32 (2023).

52. Botchway, E. et al. Resting-state network organisation in children with traumatic brain injury. Cortex 154, 89–104 (2022).

53. van der Horn, H. J. et al. Dynamic Functional Connectivity in Pediatric Mild Traumatic Brain Injury. NeuroImage 285, 120470 (2024).

54. Simchick, G. et al. Detecting functional connectivity disruptions in a translational pediatric traumatic brain injury porcine model using resting-state and task-based fMRI. Sci Rep 11, 12406 (2021).

55. Demene, C. et al. Functional ultrasound imaging of brain activity in human newborns. Science Translational Medicine 9, eaah6756 (2017).

56. Fagerholm, E. D., Hellyer, P. J., Scott, G., Leech, R. & Sharp, D. J. Disconnection of network hubs and cognitive impairment after traumatic brain injury. Brain 138, 1696–1709 (2015).

57. Manno, F. A. M. et al. Environmental enrichment leads to behavioral circadian shifts enhancing brain-wide functional connectivity between sensory cortices and eliciting increased hippocampal spiking. NeuroImage 252, 119016 (2022).

58. Rae, M. et al. Environmental enrichment enhances ethanol preference over social reward in male swiss mice: Involvement of oxytocin-dopamine interactions. Neuropharmacology 253, 109971 (2024).

59. Bellesi, G., Barker, E. D., Brown, L. & Valmaggia, L. Pediatric traumatic brain injury and antisocial behavior: are they linked? A systematic review. Brain Inj 33, 1272–1292 (2019).

60. Parker, K. J. et al. Intranasal oxytocin treatment for social deficits and biomarkers of response in children with autism. Proc Natl Acad Sci U S A 114, 8119–8124 (2017).

61. Hörnberg, H. et al. Rescue of oxytocin response and social behaviour in a mouse model of autism. Nature 584, 252–256 (2020).

62. Runyan, A., Lengel, D., Huh, J. W., Barson, J. R. & Raghupathi, R. Intranasal Administration of Oxytocin Attenuates Social Recognition Deficits and Increases Prefrontal Cortex Inhibitory Postsynaptic Currents following Traumatic Brain Injury. eNeuro 8, ENEURO.0061-21.2021 (2021).

63. Wei, Z. Z. et al. Intracranial Transplantation of Hypoxia-Preconditioned iPSC-Derived Neural Progenitor Cells Alleviates Neuropsychiatric Defects after Traumatic Brain Injury in Juvenile Rats. Cell Transplant 25, 797–809 (2016).

64. Pullela, R. et al. Traumatic Injury to the Immature Brain Results in Progressive Neuronal Loss, Hyperactivity and Delayed Cognitive Impairments. Dev Neurosci 28, 396–409 (2006).

65. Fletcher, J. L. et al. Acute treatment with TrkB agonist LM22A-4 confers neuroprotection and preserves myelin integrity in a mouse model of pediatric traumatic brain injury. Exp Neurol 339, 113652 (2021).

66. Dennis, E. L. et al. White Matter Disruption in Pediatric Traumatic Brain Injury: Results From ENIGMA Pediatric Moderate to Severe Traumatic Brain Injury. Neurology 97, e298–e309 (2021).

67. Wilde, E. A. et al. A Preliminary DTI Tractography Study of Developmental Neuroplasticity 5– 15 Years After Early Childhood Traumatic Brain Injury. Front. Neurol. 12, (2021).

68. Hsu, R.-S. et al. Wireless charging-mediated angiogenesis and nerve repair by adaptable microporous hydrogels from conductive building blocks. Nat Commun 13, 5172 (2022).

69. Ma, X. et al. Angiogenic peptide hydrogels for treatment of traumatic brain injury. Bioact Mater 5, 124–132 (2020).

70. Risen, S. R., Barber, A. D., Mostofsky, S. H. & Suskauer, S. J. Altered functional connectivity in children with mild to moderate TBI relates to motor control. J Pediatr Rehabil Med 8, 309–319 (2015).

71. Harris, J. A. et al. Hierarchical organization of cortical and thalamic connectivity. Nature 575, 195–202 (2019).

72. Scott, M. C., Bedi, S. S., Olson, S. D., Sears, C. M. & Cox, C. S. MICROGLIA AS THERAPEUTIC TARGETS AFTER NEUROLOGICAL INJURY: Strategy for Cell Therapy. Expert Opin Ther Targets 25, 365–380 (2021).

73. Favrais, G. et al. Systemic inflammation disrupts the developmental program of white matter. Ann Neurol 70, 550–565 (2011).

74. Zhang, X. et al. Cytokine toxicity to oligodendrocyte precursors is mediated by iron. Glia 52, 199–208 (2005).

75. Nemes-Baran, A. D., White, D. R. & DeSilva, T. M. Fractalkine-Dependent Microglial Pruning of Viable Oligodendrocyte Progenitor Cells Regulates Myelination. Cell Reports 32, 108047 (2020).

76. Vaughan-Jackson, A. et al. Density dependent regulation of inflammatory responses in macrophages. Front Immunol 13, 895488 (2022).

77. Hammond, T. R. et al. Single-Cell RNA Sequencing of Microglia throughout the Mouse Lifespan and in the Injured Brain Reveals Complex Cell-State Changes. Immunity 50, 253–271.e6 (2019).

78. van der Poel, M. et al. Transcriptional profiling of human microglia reveals grey-white matter heterogeneity and multiple sclerosis-associated changes. Nat Commun 10, 1139 (2019).

79. Liao, P.-Y., Chiu, Y.-M., Yu, J.-H. & Chen, S.-K. Mapping Central Projection of Oxytocin Neurons in Unmated Mice Using Cre and Alkaline Phosphatase Reporter. Front Neuroanat 14, 559402 (2020).

80. Huang, M. et al. Microglial immune regulation by epigenetic reprogramming through histone H3K27 acetylation in neuroinflammation. Front. Immunol. 14, (2023).

81. Quagliato, L. A., de Matos, U. & Nardi, A. E. Maternal immune activation generates anxiety in offspring: A translational meta-analysis. Transl Psychiatry 11, 1–6 (2021).

82. Budak, M. & Zochowski, M. Synaptic Failure Differentially Affects Pattern Formation in Heterogenous Networks. Front. Neural Circuits 13, (2019).

83. Andrew, P. M. et al. Shifts in the spatiotemporal profile of inflammatory phenotypes of innate immune cells in the rat brain following acute intoxication with the organophosphate diisopropylfluorophosphate. J Neuroinflammation 21, 285 (2024).

84. Forshammar, J. et al. Anti-inflammatory substances can influence some glial cell types but not others. Brain Research 1539, 34–40 (2013).

85. Basrai, H. S., Christie, K. J., Turbic, A., Bye, N. & Turnley, A. M. Suppressor of Cytokine Signaling-2 (SOCS2) Regulates the Microglial Response and Improves Functional Outcome after Traumatic Brain Injury in Mice. PLoS One 11, e0153418 (2016).

86. Linnerbauer, M., Wheeler, M. A. & Quintana, F. J. Astrocyte crosstalk in CNS inflammation. Neuron 108, 608–622 (2020).

87. Matejuk, A. & Ransohoff, R. M. Crosstalk Between Astrocytes and Microglia: An Overview. Frontiers in Immunology 11, (2020).

88. Witcher, K. G. et al. Traumatic brain injury-induced neuronal damage in the somatosensory cortex causes formation of rod-shaped microglia that promote astrogliosis and persistent neuroinflammation. Glia 66, 2719–2736 (2018).

89. Michelucci, A., Heurtaux, T., Grandbarbe, L., Morga, E. & Heuschling, P. Characterization of the microglial phenotype under specific pro-inflammatory and anti-inflammatory conditions: Effects of oligomeric and fibrillar amyloid-beta. J Neuroimmunol 210, 3–12 (2009).

90. Szpakowski, P., Ksiazek-Winiarek, D., Turniak-Kusy, M., Pacan, I. & Glabinski, A. Human Primary Astrocytes Differently Respond to Pro- and Anti-Inflammatory Stimuli. Biomedicines 10, 1769 (2022).

91. Da Pozzo, E. et al. Microglial Pro-Inflammatory and Anti-Inflammatory Phenotypes Are Modulated by Translocator Protein Activation. Int J Mol Sci 20, 4467 (2019).

92. Chhor, V. et al. Role of microglia in a mouse model of paediatric traumatic brain injury. Brain Behav Immun 63, 197–209 (2017).

93. Hamood, Y. et al. Sex specific effects of buprenorphine on behavior, astrocytic opioid receptor expression and neuroinflammation after pediatric traumatic brain injury in mice. Brain Behav Immun Health 22, 100469 (2022).

94. Webster, K. M. et al. Targeting high-mobility group box protein 1 (HMGB1) in pediatric traumatic brain injury: Chronic neuroinflammatory, behavioral, and epileptogenic consequences. Experimental Neurology 320, 112979 (2019).

95. Hagberg, H. et al. The role of inflammation in perinatal brain injury. Nat Rev Neurol 11, 192– 208 (2015).

96. Arambula, S. E., Reinl, E. L., El Demerdash, N., McCarthy, M. M. & Robertson, C. L. Sex differences in pediatric traumatic brain injury. Experimental Neurology 317, 168–179 (2019).

97. Schwarz, J. M., Sholar, P. W. & Bilbo, S. D. Sex differences in microglial colonization of the developing rat brain. Journal of Neurochemistry 120, 948–963 (2012).

98. Villapol, S., Loane, D. J. & Burns, M. P. Sexual dimorphism in the inflammatory response to traumatic brain injury. Glia 65, 1423–1438 (2017).

99. Tamborski, S., Mintz, E. M. & Caldwell, H. K. Sex Differences in the Embryonic Development of the Central Oxytocin System in Mice. Journal of Neuroendocrinology 28, (2016).

100. Grund, T. et al. Chemogenetic activation of oxytocin neurons: Temporal dynamics, hormonal release, and behavioral consequences. Psychoneuroendocrinology 106, 77–84 (2019).

101. Delage, C., Taib, T., Mamma, C., Lerouet, D. & Besson, V. C. Traumatic Brain Injury: An Age-Dependent View of Post-Traumatic Neuroinflammation and Its Treatment. Pharmaceutics 13, 1624 (2021).

102. Schindelin, J., et al. Fiji: an open-source platform for biological-image analysis. Nat Methods 9, 676–682 (2012).

103. Bankhead, P. et al. QuPath: Open source software for digital pathology image analysis. Sci Rep 7, 16878 (2017).

104. IJBlob: An ImageJ Library for Connected Component Analysis and Shape Analysis. Journal of Open Research Software 1, e6 (2013).

105. Ferreira, T. A. et al. Neuronal morphometry directly from bitmap images. Nat Methods 11, 982–984 (2014).

106. Effect of Aging on Elastin Functionality in Human Cerebral Arteries | Stroke. https://www.ahajournals.org/doi/10.1161/STROKEAHA.108.528091?url_ver=Z39.88-2003&rfr_id=ori:rid:crossref.org&rfr_dat=cr_pub%20%200pubmed.

107. Bokobza, C. et al. Magnetic Isolation of Microglial Cells from Neonate Mouse for Primary Cell Cultures. JoVE 62964 (2022) doi:10.3791/62964.

108. Ye, J. et al. Primer-BLAST: A tool to design target-specific primers for polymerase chain reaction. BMC Bioinformatics 13, 134 (2012).

109. Veraart, J., Fieremans, E. & Novikov, D. S. Diffusion MRI noise mapping using random matrix theory. Magn Reson Med 76, 1582–1593 (2016).

110. Veraart, J., Sijbers, J., Sunaert, S., Leemans, A. & Jeurissen, B. Weighted linear least squares estimation of diffusion MRI parameters: Strengths, limitations, and pitfalls. NeuroImage 81, 335– 346 (2013).

111. Daducci, A. et al. Accelerated Microstructure Imaging via Convex Optimization (AMICO) from diffusion MRI data. NeuroImage 105, 32–44 (2015).

112. Bonacich, P. Factoring and weighting approaches to status scores and clique identification. Journal of Mathematical Sociology (1972).

113. Noldus, L. P. J. J., Spink, A. J. & Tegelenbosch, R. A. J. EthoVision: A versatile tracking system for automation of behavioral experiments. *Behavior Research Methods*, Instruments & Computers 33, 398–414 (2001).

114. Tang, D. et al. SRplot: A free online platform for data visualization and graphing. PLoS One 18, e0294236 (2023).

