## Supplementary Figures for "Oxytocin release modulates acute neuroinflammation and improves brain development after pediatric traumatic brain injury"

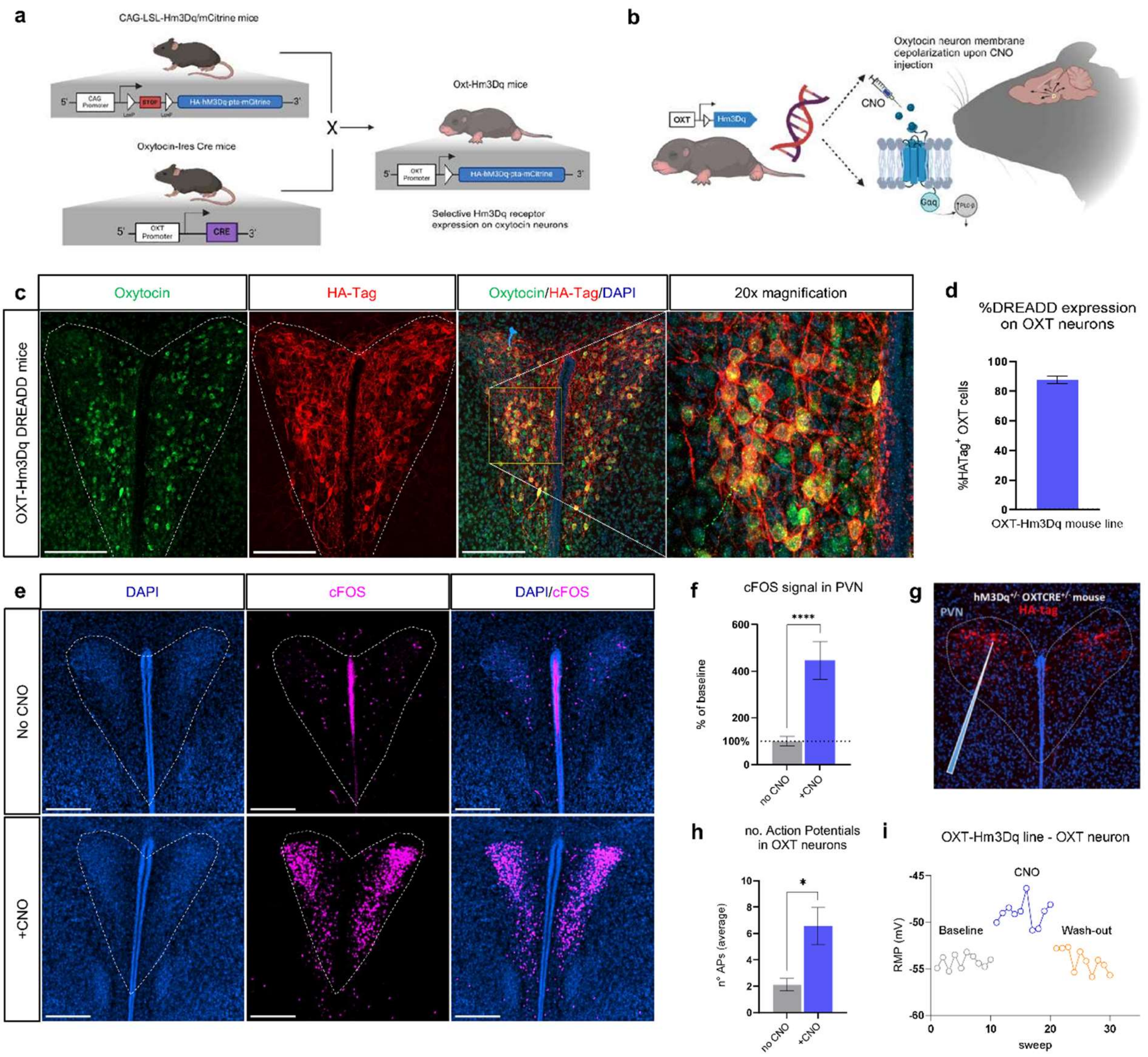


**Fig. S1: Validation of the OXT-Hm3Dq DREADD construct. a)** Graphical representation of the creation of the OXT-Hm3Dq DREADD mouse line. **b)** Graphic representing the theoretical concept of the OXT-Hm3Dq DREADD construct: Upon activation by the selective CNO ligand, designer Hm3Dq (Gq) receptors on oxytocin neurons cause membrane depolarization which creates firing of the neuron. **c)** Representative micrograph of immunohistochemistry of oxytocin (green) and HA-Tag (red) labeling, showing high coverage of Hm3Dq-DREADD receptors on oxytocin neurons (mean = 88%) in the PVN of the hypothalamus **(d).** Scale bar = 100 µm. **e)** Representative micrographs of immediate-early-gene cFOS (magenta) immunohistochemistry in the PVN in mice 1h after saline (top panel) or CNO injection (bottom panel). Scale bar = 200 µm. **f)** Signal quantification revealed a 300% increase in PVN cFOS activity in CNO-injected mice compared to the saline group. **g-i)** Electrophysiological recordings were performed on oxytocinergic neurons, identified by mCitrine fluorescence in reaction to stimulation with a 594 nm LED. Oxytocinergic neurons showed an increase in action potentials **(h)** and resting membrane potential **(i)** upon CNO administration. Based on these validation experiments, we concluded that CNO successfully causes endogenous increases in oxytocin firing, and thus formed a valid tool to execute oxytocin treatment in this study. OXT = oxytocin; DREADD = designer receptors exclusively activated by designer drugs; CNO = clozapine-n-oxide; PVN = periventricular nucleus; TBI = traumatic brain injury. In **(f)** Mann-Whitney U test (U = 0, ****$p < .0001$, no CNO: $n = 8$, +CNO: $n = 9$). In **(h)** Paired t-test ($t(4) = 2.83$, *$p = .048$, $n = 5$). Bar graphs represent Mean ± SEM.

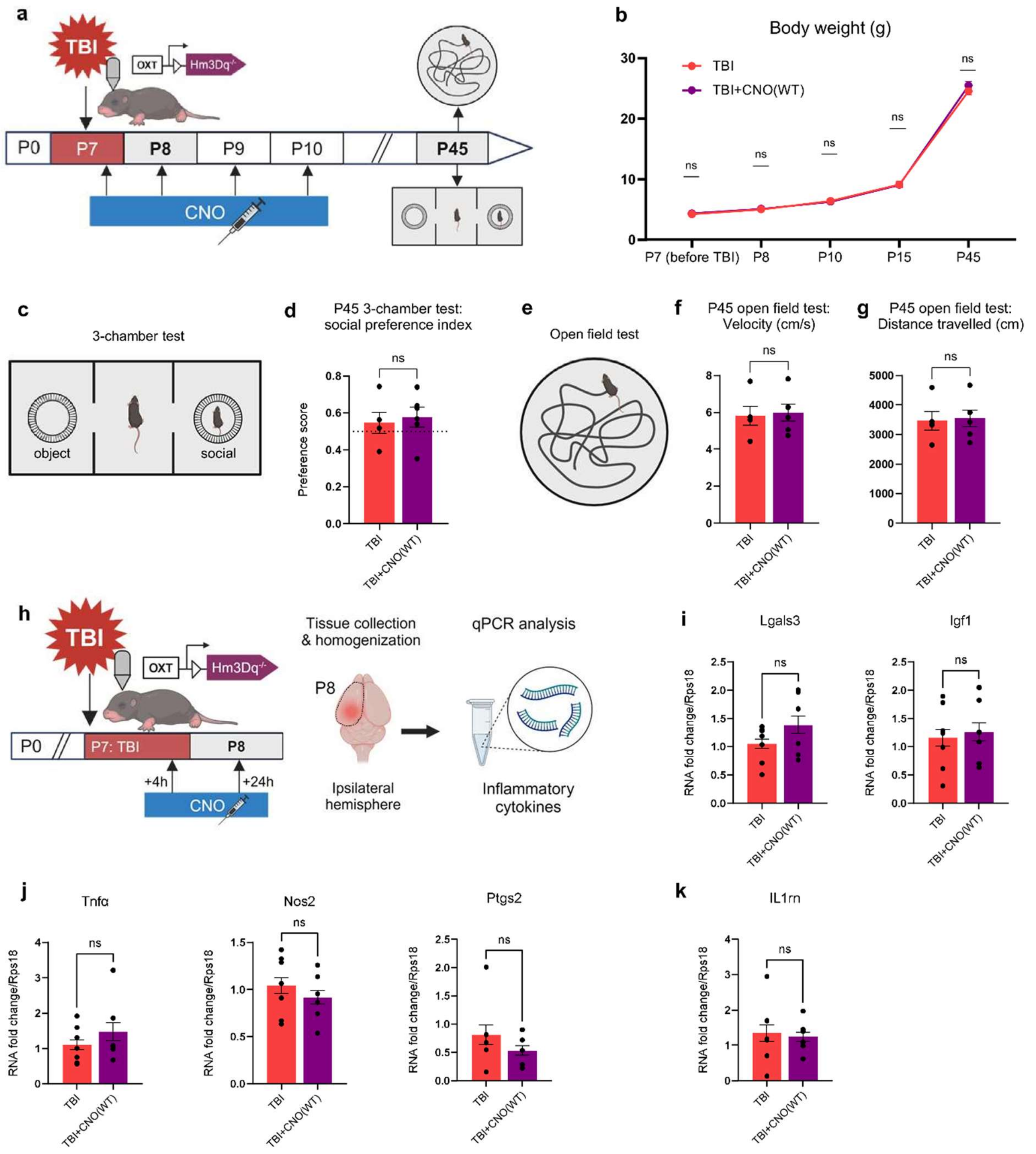


**Fig. S2: The CNO treatment paradigm of this study did not have side-effects on long-term behavior or acute neuroinflammation following TBI.** CNO in high dosages can bind to non-DREADD receptors and it can metabolize into clozapine, which is an atypical antipsychotic that can create physiological and behavioral effects in mice[1]. To assess if CNO had side-effects on readouts of our TBI model, we performed experiments on OXT-Hm3Dq$^{-/-}$ mice (which have OXT-Cre but are WT for the Hm3Dq-DREADD) subjected to TBI and treated with saline ("TBI"; red) or CNO ("TBI+CNO(WT)"; violet). **a)** Graphical timeline of behavioral experiments following P7 TBI and P7-P10 CNO treatment in WT mice. No effects of CNO were found on body weight and growth **(b)**, on social behavior during P45 3-chamber test **(c,d)**, and on activity levels during the P45 open field **(e-g)**. We performed qPCR of P8 ipsilateral neural tissue to assess the effect of CNO on neuroinflammation 24 post-injury in OXT-Hm3Dq$^{-/-}$ mice **(h)**. We did not observe a difference between TBI mice and TBI+CNO(WT) mice in anti-inflammatory **(i)**, pro-inflammatory **(j)**, and immune-regulatory **(k)** cytokine expression. These data show that the CNO treatment used in this study did not have side-effects on the acute neuroinflammatory response following TBI, nor on long-term behavior. Any therapeutic effects found by the CNO treatment in our study can thus be attributed to increased oxytocin activity, and not CNO itself. TBI = traumatic brain injury, CNO = clozapine n-oxide, WT = wild type. In **(b)** Two-way ANOVA ($F(4,36) = 1.77$, $p = 0.16$. TBI: $n = 5$, TBI+CNO(WT): $n = 6$). In **(i)** Lgals3: Mann-Whitney U test ($U = 56$, $p = 0.17$, TBI: $n = 14$, TBI+CNO(WT): $n = 12$). Igf1: Unpaired t test ($t(24) = 0.47$, $p = 0.64$, TBI: $n = 14$, TBI+CNO(WT): $n = 12$). In **(j)** Tnfα: Mann-Whitney U test ($U = 64$, $p = 0.33$, TBI: $n = 14$, TBI+CNO(WT): $n = 12$). Nos2: Unpaired t test ($t(24) = 1.16$, $p = 0.26$, TBI: $n = 14$, TBI+CNO(WT): $n = 12$). Ptgs2 : Mann-Whitney U test ($U = 48$, $p = 0.47$, TBI: $n = 12$, TBI+CNO(WT): $n = 10$). In **(k)** Unpaired t test ($t(24) = 0.39$, $p = .70$, TBI: $n = 14$, TBI+CNO(WT): $n = 12$). Bar graphs represent Mean ± SEM. Each circle corresponds to one animal.

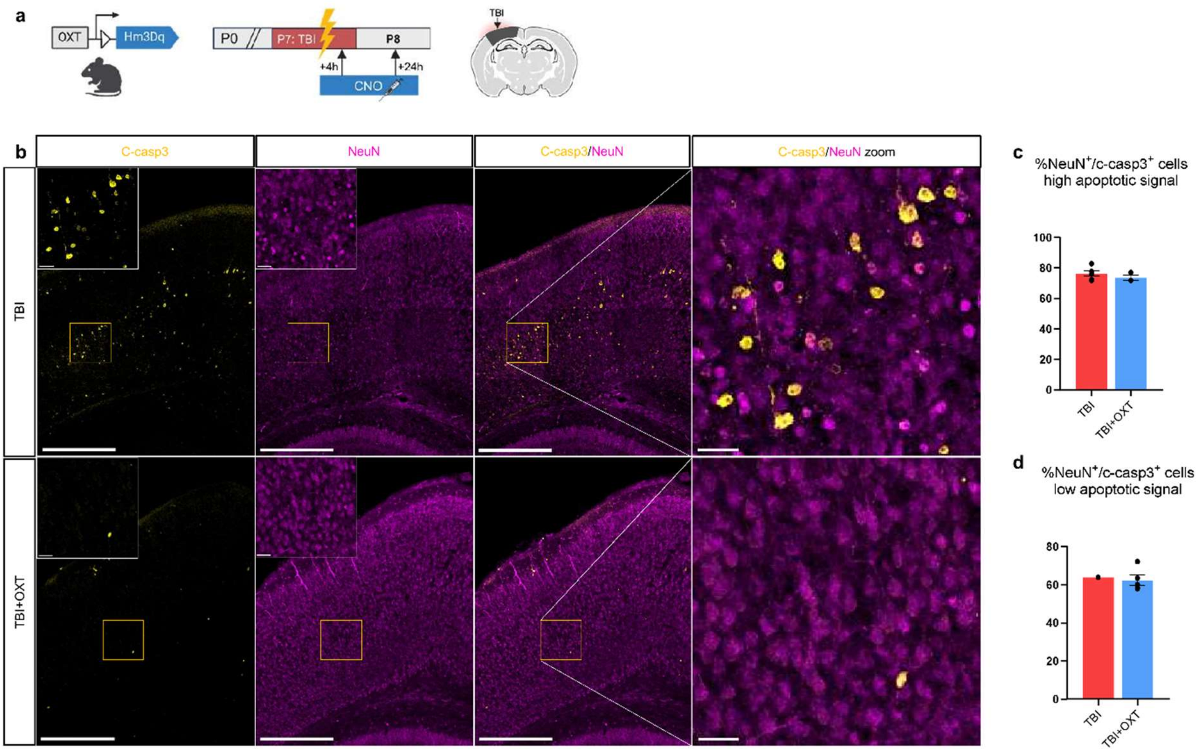


**Fig. S3: Oxytocin does not affect the proportion neuronal apoptotic death following TBI. a)** Graphical timeline of P8 immunohistochemistry experiment after P7 TBI and 2 sessions of CNO treatment, and graphical indication of the ipsilateral somatosensory cortex ROI. **b)** Representative micrographs of apoptotic cell death (cleaved-caspase 3; yellow) and neuron (NeuN; magenta) immunoreactivity in TBI (top row) and TBI+OXT (bottom row) mice. Scale bar = 200 µm (inserts: scale bar = 30 µm). To eliminate confounding effects of apoptotic signal degree on the proportion neuronal cell death, quantification was divided into high apoptotic and low apoptotic signal. **c,d)** Quantification of the proportion of neuronal cell death in samples with **(c)** high apoptotic and **(d)** low apoptotic reactivity showed no difference between TBI and TBI+OXT mice. TBI = traumatic brain injury. CNO = clozapine n-oxide, ROI = region of interest, c-casp3 = cleaved-caspase 3, OXT = oxytocin. In **(c)** Unpaired t-test ($t(7) = 1.04$, $p = .33$, TBI: $n = 6$, TBI+OXT: $n = 3$). In **(d)** sample size too small to perform statistical testing (TBI: $n = 1$, TBI+OXT: $n = 5$). Bar graphs represent Mean ± SEM. Each circle represents an individual sample.

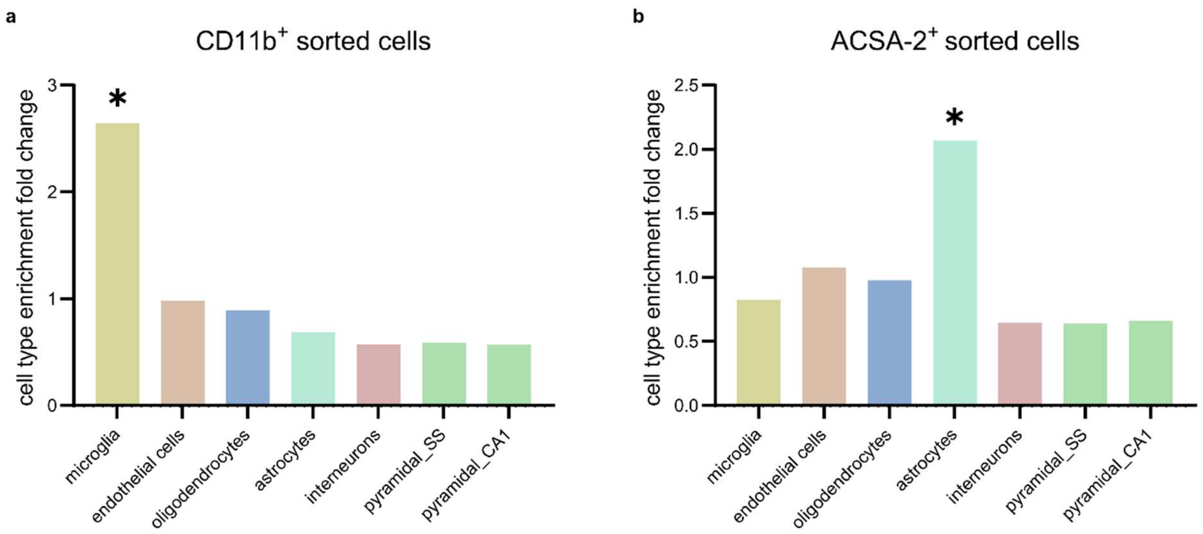


**Fig S4: Cell type enrichment analysis confirms purity of magnetically sorted microglia and astrocytes.** MACS® (Miltenyi Biotech) magnetic sorting purity of CD11b$^+$ and ACSA-2$^+$ cells was assessed with Expression Weighted Cell-type Enrichment (EWCE) analysis[2] based on top 250 expressed genes in the respective RNA Seq datasets. **a,b)** Bar graphs showing enrichment fold change of neural cell type marker gene expression in our transcriptomic datasets, including microglia, endothelial cells, oligodendrocytes, astrocytes, interneurons and pyramidal cells. Asterisks represent cell type enrichment with $p < .05$. **a)** EWCE analysis revealed a significant enrichment of only microglia cells in the CD11b$^+$ transcriptomic dataset. **b)** EWCE analysis revealed a significant enrichment of only astrocytes in the ACSA-2$^+$ transcriptomic dataset. This analysis confirms that the CD11b$^+$ and ACSA-2$^+$ sorted cells were indeed microglia and astrocytes, respectively.

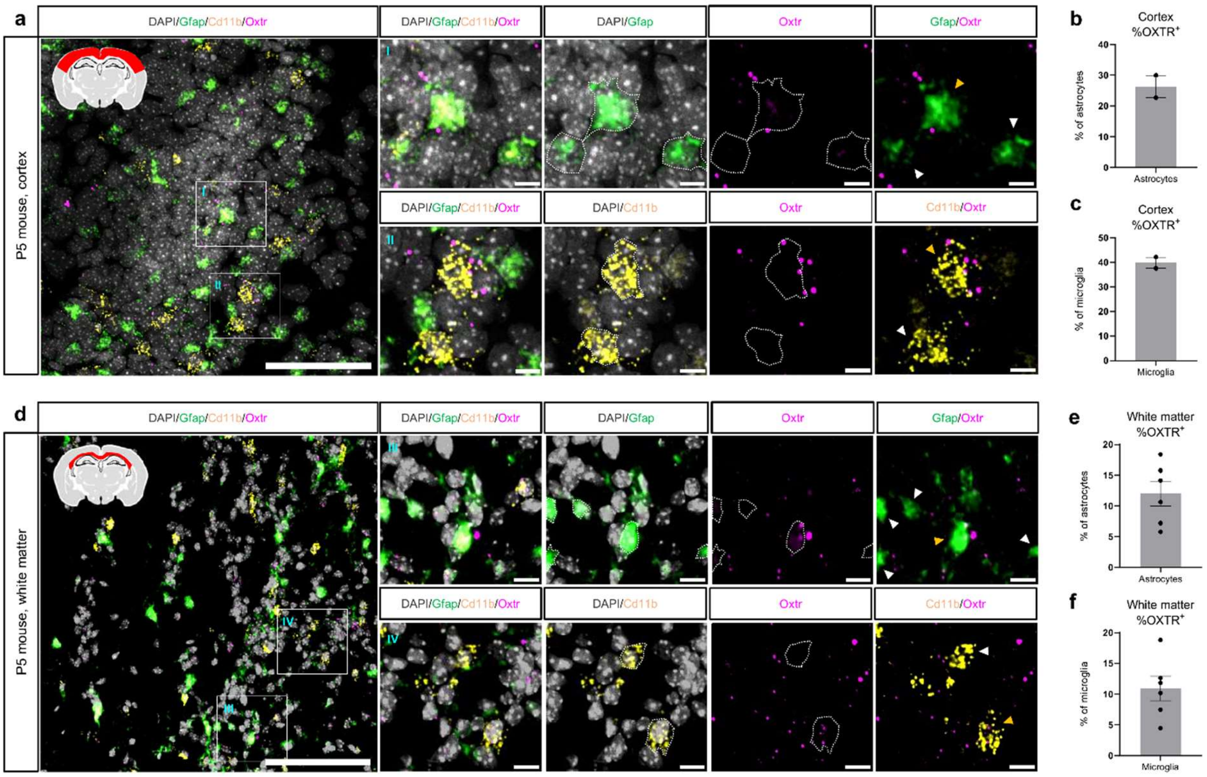


**Fig. S5: RNA Scope in situ hybridization reveals OXT receptor expression on microglia and astrocytes in P5 mice.** To assess if microglia and astrocytes express oxytocin receptors (OXTR) *in-vivo* in mouse pups, we performed RNA Scope in situ hybridization on brain sections from untreated P5 OXT-Hm3Dq mice. **a)** ROI indication (red) and representative micrograph from cortex showing reactivity of oxytocin receptor (*Oxtr*; magenta), microglia (*CD11b*; yellow), and astrocyte (*Gfap*; green) mRNA. Scale bar = 100 µm. Inserts highlight cortical astrocytes (top row) and microglia (bottom row) with (yellow arrow) and without (white arrow) expression of *Oxtr* mRNA. Scale bar = 10 µm. **b,c)** Quantification of micrographs revealed that 26% of cortical astrocytes **(b)** and 39% of cortical microglia express OXTR **(c)**. **d)** ROI indication (red) and representative micrograph from white matter showing *Oxtr*, *Gfap* and *CD11b* mRNA reactivity in the white matter. Scale bar = 100 µm. Inserts highlight white matter astrocytes (top row) and microglia (bottom row) with (yellow arrow) and without (white arrow) expression of OXTR mRNA. Scale bar = 10 µm. **e,f)** Quantification of micrographs revealed that 12% of white matter astrocytes **(e)** and 11% of white matter microglia express OXTR **(f)**. These data confirm oxytocin receptor expression on microglia and astrocytes, suggesting both cell types can be directly influenced by neuronal oxytocin. Bar graphs represent Mean ± SEM. Each circle represents an individual sample.

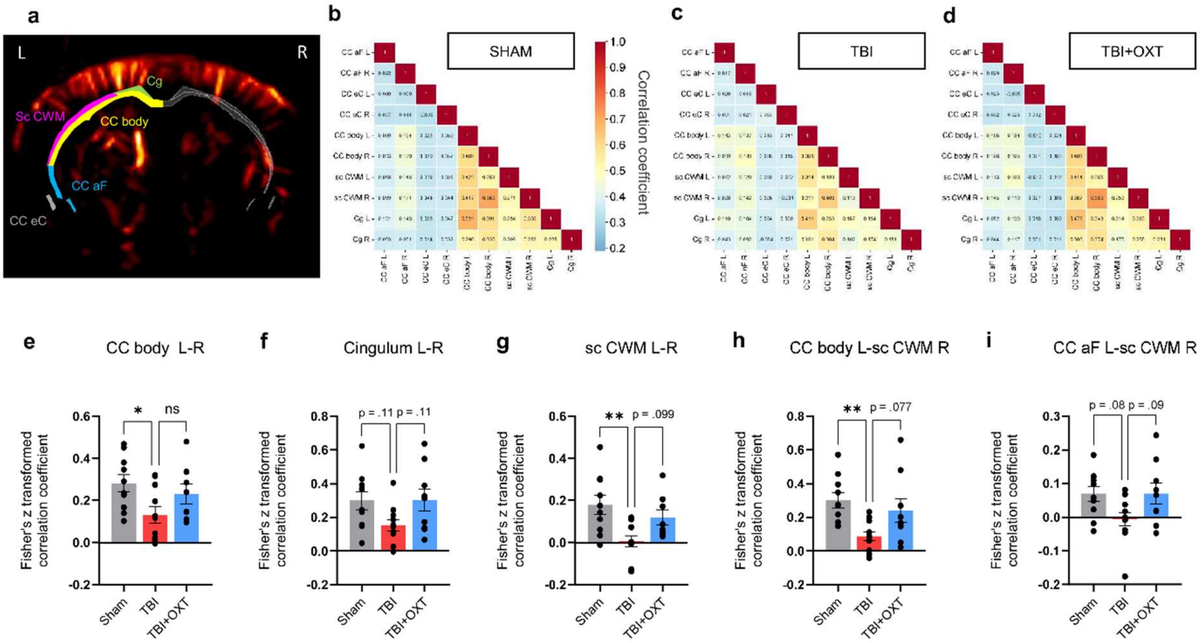


**Fig. S6: Oxytocin affects long-term functional connectivity of white matter regions following TBI. a)** Representative image of functional ultrasound (fUS) signal and the white matter ROIs in P45 mice: Cg (green), CC body (yellow), sc CWM (magenta), CC aF (blue), CC eC (gray). **b-d)** Average correlation coefficient matrices of functional connectivity between the white matter ROIs in the Sham **(b)**, TBI **(c)** and TBI+OXT **(d)** groups. **e-i)** Quantification of correlation coefficients between a selection of white matter ROI pairs, showing reduced functional connectivity in TBI, and reversed functional connectivity after oxytocin treatment. TBI = traumatic brain injury; CNO = clozapine n-oxide; fUS = functional ultrasound; ROI = region of interest; Cg = cingulum, CC body = corpus callosum body, sc CWM = supra-callosal cerebral white matter, CC aF = corpus callosum anterior forceps, CC eC = corpus callosum external capsule. In **(e)** ANOVA (F(2,26) = 3.69, *$p$ = .039, Sham: $n$ = 10, TBI: $n$ = 11, TBI+CNO: $n$ = 8). Tukey's (Sham vs. TBI: *$p$ = .034; Sham vs. TBI+OXT: $p$ = .69; TBI vs. TBI+OXT: $p$ = .24). In **(f)** ANOVA (F(2,27) = 2.98, $p$ = .068, Sham: $n$ = 10, TBI: $n$ = 11, TBI+CNO: $n$ = 9). Tukey's (Sham vs. TBI: $p$ = .11; Sham vs. TBI+OXT: $p$ = 1; TBI vs. TBI+OXT: $p$ = .11). In **(g)** ANOVA (F(2,26) = 6.32, **$p$ = .006, Sham: $n$ = 10, TBI: $n$ = 11, TBI+CNO: $n$ = 8). Tukey's (Sham vs. TBI: **$p$ = .005; Sham vs. TBI+OXT: $p$ = .52; TBI vs. TBI+OXT: $p$ = .099). In **(h)** ANOVA (F(2,27) = 5.69, **$p$ = .009, Sham: $n$ = 10, TBI: $n$ = 11, TBI+CNO: $n$ = 9). Tukey's (Sham vs. TBI: **$p$ = .008; Sham vs. TBI+OXT: $p$ = .66; TBI vs. TBI+OXT: $p$ = .077). In **(i)** ANOVA (F(2,27) = 3.44, *$p$ = .047, Sham: $n$ = 10, TBI: $n$ = 11, TBI+CNO: $n$ = 9). Tukey's (Sham vs. TBI: $p$ = .077; Sham vs. TBI+OXT: $p$ = 1; TBI vs. TBI+OXT: $p$ = .089). Bar graphs represent Mean ± SEM. Each circle represents an individual sample.

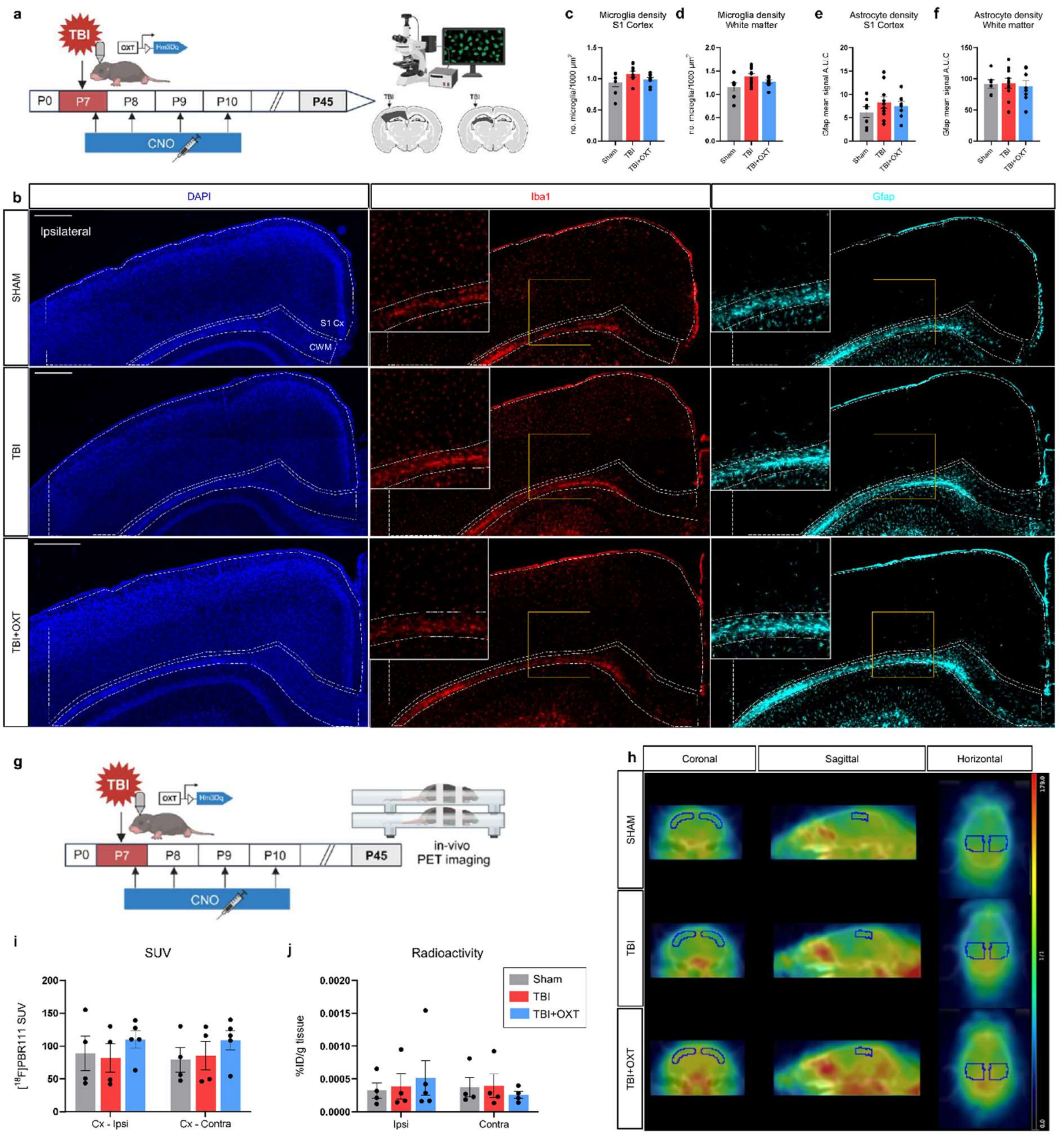


**Fig. S7: Early-life TBI is not associated with long-term neuroinflammation. a)** Graphical timeline of the P45 immunohistochemistry experiment with cortical and WM ROI (black) indication. **b)** Representative micrograph of Iba1$^+$ microglia (red) and GFAP$^+$ astrocytes (cyan) in ipsilateral S1 (dotted lines, top) and ipsilateral WM (dotted lines, bottom) for Sham, TBI and TBI+OXT groups. Scale bar = 400 µm. **c,d)** Quantification of number of microglia cells per 100 µm$^2$ in ipsilateral S1 **(c)**, and ipsilateral WM **(d)**, revealed no significant difference in microglia density between experimental groups at P45. **e,f)** Quantification of GFAP signal density in ipsilateral S1 **(e)**, and ipsilateral WM **(f)**, revealed no difference in astrocyte density between experimental groups at P45. **g)** Graphical timeline of P45 TSPO assessment using *in-vivo* positron emission tomography (PET) imaging and *ex-vivo* gamma counting. **h)** Average SUV images of [$^{18}$F]PBR111 acquired by PET imaging in sham, TBI and TBI+OXT animals, co-registered to the CT in the coronal (left), sagittal (center) and horizontal (right) planes. Blue areas correspond to volumes of interest (VOIs). **i)** Quantification of [$^{18}$F]PBR111 SUV in ipsilateral and contralateral VOI in the three experimental groups. TSPO expression is not modulated by TBI or OXT treatment. **j)** Quantification of radioactivity (% injected dose (ID)/g of tissue) in the ipsilateral and contralateral VOI. Results confirm the absence of TSPO overexpression after TBI at long term. ROI = region of interest; S1 = primary somatosensory cortex; CWM = cingulate white matter; TBI = traumatic brain injury; OXT = oxytocin; TSPO = 18 kDa translocator protein; SUV = standard uptake value; VOI = volume of interest. In **(c)** ANOVA (F(2,22) = 1.80, *p* = .19, Sham: *n* = 7, TBI: *n* = 10, TBI+CNO: *n* = 8). In **(d)** ANOVA (F(2,22) = 3.16, *p* = .06, Sham: *n* = 7, TBI: *n* = 10, TBI+CNO: *n* = 8). In **(e)** ANOVA (F(2,22) = 0.87, *p* = .43, Sham: *n* = 7, TBI: *n* = 10, TBI+CNO: *n* = 8). In **(f)** ANOVA (F(2,21) = 0.11, *p* = .90, Sham: *n* = 6, TBI: *n* = 10, TBI+CNO: *n* = 8). In **(i)** Two-way ANOVA (F(2,20) = 0.06, *p* = .94, Sham: *n* = 4, TBI: *n* = 5, TBI+OXT: *n* = 5). In **(j)** Two-way ANOVA (F(2,19) = 0.42, *p* = 0.66. Sham: *n* = 4, TBI: *n* = 5, TBI+CNO: *n* = 5). Bar graphs represent Mean ± SEM. Each circle corresponds to one animal.

**References**


1. Manvich, D. F. *et al.* The DREADD agonist clozapine N-oxide (CNO) is reverse-metabolized to clozapine and produces clozapine-like interoceptive stimulus effects in rats and mice. *Sci Rep* **8**, 3840 (2018).
2. Skene, N. G. & Grant, S. G. N. Identification of Vulnerable Cell Types in Major Brain Disorders Using Single Cell Transcriptomes and Expression Weighted Cell Type Enrichment. *Front. Neurosci.* **10**, (2016).
