## Supplementary Methods for "Oxytocin release modulates acute neuroinflammation and improves brain development after pediatric traumatic brain injury"

### A Supplementary experiments to validate the OXT-Hm3Dq DREADD construct

#### A.1 Immunohistochemistry

##### A.1.1 Hm3Dq DREADD expression on oxytocin neurons

Validation immunohistochemistry of the OXT-Hm3Dq mouse line was performed to assess the degree of DREADD expression on oxytocin cells in the periventricular nucleus (PVN) – the principal production region of oxytocin[1]. Untreated OXT-Hm3Dq mice were anesthetized and sacrificed as described in the main methods. 50 µm free-floating coronal sections were cut with a vibratome. The immunohistochemistry was a 5-day experiment. Day 1: Sections were incubated for 1h in blocking solution (10%BSA + 3% NGS in PBS1x), and incubated at 4°C for 48h with mouse anti-HA-Tag (cell signaling 2367; 1:200) in incubation solution (3%BSA + 1%NGS+0.2%TritonX in PBS1x). Day 3: Sections were washed in 0.2%TritonX+PBS1x and incubated at RT for 2h with goat anti-mouse Alexa 594 (Abcam ab150116; 1:500). Sections were washed in 0.1%TritonX in PBS1x, blocked for 45 min in blocking solution and subsequently incubated at 4°C for 48h with rabbit anti oxytocin (Millipore Ab911; 1:500) in incubation solution. Day 5: Sections were washed in 0.2%TritonX+PBS1x and incubated at RT for 2h with goat anti rabbit Alexa 488 (Abcam ab150077; 1:500). Slides were washed, mounted and coverslipped with Fluoromount (Abcam ab104139).

##### A.1.2 c-FOS activation upon CNO injection

We validated the increase in PVN cell firing after CNO injection with immunohistochemistry of immediate-early-gene c-FOS[2]. OXT-Hm3Dq mice were injected with saline or CNO solution, and sacrificed under anesthesia 1h post-injection. Brains were collected and cut into 50 µm floating sections with a vibratome. Sections were blocked for 1h in 10% BSA + 3%NGS + 0.2% TritonX in PBS 1x, and incubated at 4°C for 24h with guinea pig anti-c-Fos (synaptic systems 226 308; 1:5000) in 3% BSA + 1%NGS + 0.2% TritonX in PBS1x. Sections were washed in PBS1x and incubated with sheep anti guinea pig FITC (Abcam ab136790; 1:500) in PBS1x + 0.1% TritonX for 2h at RT. Sections were washed, mounted and coverslipped with Fluoromount (Abcam ab104139).

##### A.1.3 Image acquisition

Brain sections were imaged with a ZEISS LSM 800 Airyscan confocal microscope (Zeiss, Oberkochen, Germany). Images of the PVN were acquired as 3x4 tiles of 20x z-stack images with 0.49 µm step size.

##### A.1.4 Image analysis

Image analysis was performed with FIJI/ImageJ[3] via manual counting by researchers blinded to experimental conditions. The degree of HA-Tag coverage of OXT cells in the PVN was determined as the number of HATag$^{+}$/OXT$^{+}$ cells divided by the total number of OXT$^{+}$ cells. For the cFOS quantification, cFOS$^{+}$ cells were counted and transformed as a percentage relative to the average cFOS signal of the saline injected group ("% of baseline").

A.2 Electrophysiology

Coronal sections of the midbrain, 250 µm thick and including the PVN, were obtained from mice of the OXT-Hm3Dq DREADD line. Brain slicing was performed using a cutting solution composed of 90.89 mM choline chloride, 24.98 mM glucose, 25 mM NaHCO3, 6.98 mM MgCl2, 11.85 mM ascorbic acid, 3.09 mM sodium pyruvate, 2.49 mM KCl, 1.25 mM NaH2PO4, and 0.50 mM CaCl2. The sections were maintained in the same solution for 20–30 minutes at 35°C. Subsequently, the slices were transferred to artificial cerebrospinal fluid (aCSF) composed of 119 mM NaCl, 2.5 mM KCl, 1.3 mM MgCl2, 2.5 mM CaCl2, 1.0 mM NaH2PO4, 26.2 mM NaHCO3, and 11 mM glucose, continuously bubbled with a gas mixture of 95% O2 and 5% CO2 at room temperature. Electrophysiological recordings, in either whole-cell voltage-clamp or current-clamp mode, were performed at 35–37°C with slices submerged in aCSF perfused at 2–3 ml/min. The internal solution for recording pipettes consisted of 140 mM K-Gluconate, 2 mM MgCl2, 5 mM KCl, 0.2 mM EGTA, 10 mM HEPES, 4 mM Na2ATP, 0.3 mM Na3GTP, and 10 mM creatine-phosphate. Oxytocinergic neurons were identified by stimulating the tissue with a 594 nm LED to detect mCitrine fluorescence. The resting membrane potential (measured in mV) was recorded using a Multiclamp 700B Commander (Molecular Devices) under zero current injection conditions (I=0) immediately after establishing whole-cell configuration. Action potentials were induced in current-clamp mode by applying depolarizing current steps of 100 pA lasting 500 ms. Following baseline acquisition, CNO (20 µM, Enzo Life Sciences, BML-NS-105-0025) was administered to the bath solution and subsequently washed out. Throughout the experiment, the number of action potentials and resting membrane potential values were systematically monitored.

### B Supplementary experiments to assess side-effects of the CNO substance

B.1 Animals

Cre-dependent Hm3Dq-DREADD (Designer Receptors Exclusively Activated by Designer Drugs) mice (B6N;129-Tg(CAG-CHRM3*,-mCitrine)1Ute/J[4], Jackson stock #026220) were crossed with Oxytocin-Ires-Cre mice (B6;129S-Oxttm1.1(cre)Dolsn/J, Jackson stock #024234) to create OXT$^{+/-}$-Hm3Dq$^{-/-}$ mice. We used these mice to test side-effects of the CNO substance because they come from the same breeding colony and they share the exact same genetic background as the OXT-Hm3Dq experimental mice, but without DREADD expression. As such, the CNO substance could be tested isolated from its functional effects on the Gq designer receptors.

TBI injury and CNO treatment were administered to the $OXT^{+/-}$-$Hm3Dq^{-/-}$ animals as described in the main methods.

B.2 real-time qPCR

RNA was extracted from P8 ipsilateral neural tissue using the Nucleospin RNA XS extraction kit (Macherey-Nagel). RNA quantity was measured with Nanodrop 2000 and was normalized across samples. RNA was subjected to reverse transcriptase using the MLV-RT kit (Promega; M1701). Real-time quantitative transcription polymerase chain reaction (RT-qPCR) was performed in duplicates with PowerUp™ SYBR™ Green Master Mix (ThermoFischer) for 40 cycles with a 2-step program (15 sec of denaturation at 95°C and 1 min of annealing/extension at 60°C). mRNA amplification values were normalized on Rps18 housekeeping gene and calculated as mRNA fold change measures normalized on the TBI+saline group following the ΔΔCt analysis method. We tested a selection of microglia-related inflammation genes, based on previous findings[5]: Nos2, Ptgs2, Tnf-α, Lgals3, Igf1, and IL1rn. Primers were designed with Primer-BLAST [6]. Primer sequences are provided in Table S2.

B.3 Behavior

To assess long-term side-effects of CNO on behavior, we subjected the CNO-treated and saline-treated $OXT^{+/-}$-$Hm3Dq^{-/-}$ TBI mice to P45 3-chamber test and P45 open field test. Data acquisition and analysis was performed as described in the main methods.

<u>C Supplementary experiment on *in-vivo* oxytocin receptor mRNA expression</u>

To assess oxytocin receptor (OXTR) expression on microglia and astrocytes *in-vivo* in young mice (P5), we performed RNA Scope in situ hybridization of Oxtr mRNA expression.

C.1 Fluorescent RNAscope in situ hybridization

RNA in situ hybridization was performed using RNAScope® Multiplex Fluorescent V2 Assay with the ACD HybEZ™ II Hybridization System (Advanced Cell Diagnostics, Hayward, California, USA), according to the manufacturer's protocol. Mouse-specific probes for oxytocin receptor (RNAscope® Probe #412171), Cd11b (RNAscope® Probe #311491-C2) and Gfap (RNAscope® Probe #313211-C3) mRNA were used to identify OXTR expression on microglia and astrocytes, respectively. Probes were amplified and fluorescently labeled using TSA Vivid Fluorofore kits 520 (#7523), 570 (#7526) and 650 (#7527). Slides were coverslipped with DAPI-immersed mounting medium to visualize cell nuclei.

C.2 Image acquisition

Fluorescent brain sections were imaged with a ZEISS LSM 800 Airyscan confocal microscope (Zeiss, Oberkochen, Germany) at 20x magnitude. 5 separate images were taken in the cortex and white matter, for each animal.

C.3 Image analysis

Images analysis was performed with FIJI/ImageJ[3]. Astrocytes and microglia cells were counted based on co-labeling of DAPI cell nuclei with Gfap and Cd11b mRNA, respectively. The degree of $OXTR^+$ astrocytes/microglia was calculated as the number of $OXTR^+$ astrocytes/microglia divided by the total number of astrocyte/microglia cells. Data from 5 images was averaged for each sample. Quantification was performed separately for the cortex and white matter.

D Supplementary experiment on acute neuronal cell death

To assess the proportion neuronal cell death after TBI and oxytocin exposure, we performed immunohistochemistry on P8 mouse brains from TBI and TBI+OXT groups.

D.1 Immunohistochemistry

Immunohistochemistry was performed on P8 brain sections as described in the main methods. Primary antibodies used were mouse anti-NeuN (Sigma-Aldrich MAB377; 1:200) and rabbit anti-cleaved-caspase 3 (Cell Signaling D175; 1:200). Secondary antibodies used were donkey anti mouse 555 (Abcam ab150106; 1:500) and donkey anti rabbit 488 (Abcam ab150061; 1:500).

D.2 Image acquisition

Brain sections were imaged with a ZEISS Axioscan.Z1 (Zeiss, Oberkochen, Germany) at 10x.

D.3 Image analysis

Image analysis was performed by researchers blinded to experimental conditions. NeuN and cleaved-caspase 3 signals were quantified in the cortex ipsilateral to the brain injury, by manual counting. The degree of neuronal apoptotic cell death was quantified per animal as the number of $NeuN^+$/cleaved-caspase-$3^+$ cells divided by the total number of cleaved-caspase $3^+$ cells.

E Supplementary experiments on long-term neuroinflammation after TBI

To assess the long-term neuroinflammatory profile of our TBI model, we performed P45 experiments of microglia and astrocyte reactivity in sham, TBI and TBI+OXT groups.

E.1 P45 Immunohistochemistry

Immunohistochemistry of Iba1$^+$ (microglia) and Gfap$^+$ (astrocyte) reactivity was performed on free-floating sections from P45 brains as described in the main methods. Microglia and astrocyte density were analyzed in the ipsilateral somatosensory cortex and white matter as described in the main methods. Analysis was performed by a researcher blinded to experimental conditions. Microglia were manually counted, and astrocyte density was analyzed per sample as mean signal intensity measures corrected for background signal.

E.2 P45 TSPO quantification

To complement the microgliosis assay, we set out to measure the presence of 18 kDa Translocator Proteins (TSPO) at P45, which are proteins expressed by glia, that can act as a biomarker of neuroinflammation (reviewed in Cumbers et al., 2024[7]).

E.2.1 [$^{18}$F]PBR111 PET imaging and processing

P45 animals were anaesthetised with isoflurane (4% induction, maintained at 2%) and injected with [$^{18}$F]PBR111 in the tail vein immediately prior to imaging – [$^{18}$F]PBR111 is a second-generation PET ligand that specifically binds the TSPO. To maintain the same specific activity between animals at the time of injection, all syringes were prepared at the same time, the injected dose consequently varied from 33.54 MBq to 5.76 MBq. A CT image was acquired, followed by a dynamic PET acquisition of 50 minutes per animal (5x10 min), during which body temperature was maintained using a heated bed. Images were acquired on a FLEX TriumphTM preclinical PET-CT scanner (Gamma Medica-Ideas, Nortridge, CA).

[$^{18}$F]PBR111 dynamic images were co-registered to a T2-weighted MRI atlas using PMOD (v4.401, PMOD Technologies LLC). A factorial analysis was applied to extract the signal component attributed to defluorination-related accumulation in the skull, using PET data processed with Pixies (Apteryx), as previously described[8]. Static images were then created for the last 15 min of scan. Standard uptake values (SUV) corrected for injected dose and weight were then generated using Statistical Parametric Mapping (SPM12) with the Small Animal Molecular Imaging Toolbox (SAMIT; v3.0). A volume of interest (VOI) was then manually created in the TBI-targeted area and analysed in both hemispheres using PMOD.

E.2.2 *Ex-vivo* radioactivity counting

Radioactive concentrations in targeted TBI area and the contralateral area were measured on an automatic γ counting system (Wizard 3", PerkinElmer). Data are expressed in percentage of the injected dose (% ID)/g of tissue for each animal.

**References**

1. Matsunaga, W. *et al.* LPS-induced Fos expression in oxytocin and vasopressin neurons of the rat hypothalamus. *Brain Res* **858**, 9–18 (2000).
2. Hoffman, G. E., Smith, M. S. & Verbalis, J. G. c-Fos and related immediate early gene products as markers of activity in neuroendocrine systems. *Front Neuroendocrinol* **14**, 173–213 (1993).
3. Schindelin, J. *et al.* Fiji: an open-source platform for biological-image analysis. *Nat Methods* **9**, 676–682 (2012).
4. Zhu, H. *et al.* Cre dependent DREADD (Designer Receptors Exclusively Activated by Designer Drugs) mice. *Genesis (New York, N.Y. : 2000)* **54**, 439 (2016).
5. Jacquens, A. *et al.* Deleterious effect of sustained neuroinflammation in pediatric traumatic brain injury. *Brain, Behavior, and Immunity* **120**, 99–116 (2024).
6. Ye, J. *et al.* Primer-BLAST: A tool to design target-specific primers for polymerase chain reaction. *BMC Bioinformatics* **13**, 134 (2012).
7. Cumbers, G. A., Harvey-Latham, E. D., Kassiou, M., Werry, E. L. & Danon, J. J. Emerging TSPO-PET Radiotracers for Imaging Neuroinflammation: A Critical Analysis. *Seminars in Nuclear Medicine* **54**, 856–874 (2024).
8. Millet, P. *et al.* Quantification of dopamine D2/3 receptors in rat brain using factor analysis corrected [18F]Fallypride images. *NeuroImage* **62**, 1455–1468 (2012).
